# Extremely widespread parthenogenesis and a trade-off between alternative forms of reproduction in mayflies (Ephemeroptera)

**DOI:** 10.1101/841122

**Authors:** Maud Liegeois, Michel Sartori, Tanja Schwander

## Abstract

Studying alternative forms of reproduction in natural populations is of fundamental importance for understanding the costs and benefits of sex. Mayflies are one of the few animal groups where sexual reproduction co-occurs with different types of parthenogenesis, providing ideal conditions for identifying benefits of sex in natural populations. Here, we establish a catalogue of all known mayfly species capable of reproducing by parthenogenesis, as well as mayfly species unable to do so. Overall, 1.8% of the described species reproduce parthenogenetically, which is an order of magnitude higher than reported in other animal groups. This frequency even reaches 47.8% if estimates are based on the number of studied rather than described mayfly species. In terms of egg-hatching success, sex is a more successful strategy than parthenogenesis, and we found a trade-off between the efficiency of sexual and parthenogenetic reproduction across species. This means that improving the capacity for parthenogenesis may come at the cost of being less able to reproduce sexually, even in facultative parthenogens. Such a trade-off can help explain why facultative parthenogenesis is extremely rare among animals despite its potential to combine the benefits of sexual and parthenogenetic reproduction. We argue that parthenogenesis is frequently selected in mayflies in spite of this probable trade-off because their typically low dispersal ability and short and fragile adult life may frequently generate situations of mate limitation in females. Mayflies are currently clearly underappreciated for understanding the benefits of sex under natural conditions.

## INTRODUCTION

The evolution and maintenance of sexual reproduction has been one of the major questions in evolutionary ecology for the last decades (e.g., Agrawal, 2006; Otto, 2009; Jalvingh *et al.*, 2016). Sex is associated with profound costs (reviewed in Lehtonen *et al.*, 2012), yet it is the most widespread reproductive mode among animals. Female-producing parthenogenesis (thelytoky) would largely avoid the costs associated with sex, yet only a minority of animals are known to reproduce parthenogenetically. Whether this minority is small or rather just slim remains however unknown as there are only two quantitative estimates (based on species lists) of the frequency of parthenogenesis (i.e., in vertebrates: White, 1973; Vrijenhoek, 1998; and in haplodiploids: van der Kooi *et al.*, 2017). This is unfortunate as such species lists are invaluable for addressing when and how parthenogenetic reproduction is favoured over sex in natural populations (e.g., Ross *et al.*, 2013; van der Kooi *et al.*, 2017) and thus for helping to solve the paradox of sex. In this review, we provide such a quantitative estimate by summarising the current knowledge on sexual and parthenogenetic reproduction in Ephemeroptera (mayflies), as a first step towards developing this group for the study of benefits of sex in natural populations.

Mayflies constitute one of the most basal (early diverging) lineage of insects (Edmunds and McCafferty, 1988), their origin dating back to ~300 Mya (Brittain and Sartori, 2009). Widespread around the world with 3’666 described species (42 families, 472 genera; adapted from Sartori and Brittain, 2015; MS pers. com.), they are well studied for being an important link in the food chain, for their use for fly fishing (Knopp and Cormier, 1997), and for their potential as bioindicators of water quality (Bauernfeind and Moog, 2000). Mayflies do not feed as adults, relying solely on the reserves accumulated during their aquatic larval stages. Adult life span is extremely short, lasting from few hours to some days depending on the species. Because of their typically low dispersal ability and their short and fragile adult life, mayflies have restricted opportunities for reproduction, which we argue is one of the factors that may have selected for the evolution of parthenogenesis in this group. Parthenogenesis in mayflies can be largely accidental (i.e., tychoparthenogenesis), facultative or ‘obligate’ (see Box 1). Furthermore, a single species can feature mixed reproduction (some females reproduce sexually, others parthenogenetially), either sympatrically or in allopatry (i.e., geographical parthenogenesis).

We conducted a detailed literature review to establish a catalogue of all (to the best of our knowledge) mayfly species capable of reproducing parthenogenetically, and study whether the frequency of parthenogenesis varies among mayfly clades. We then use this catalogue to conduct cross-species comparisons with respect to the cellular mechanisms of parthenogenesis, how the capacity for parthenogenesis affects population sex ratios, as well as the geographical distribution of sexual and parthenogenetic populations.

**Box 1.**
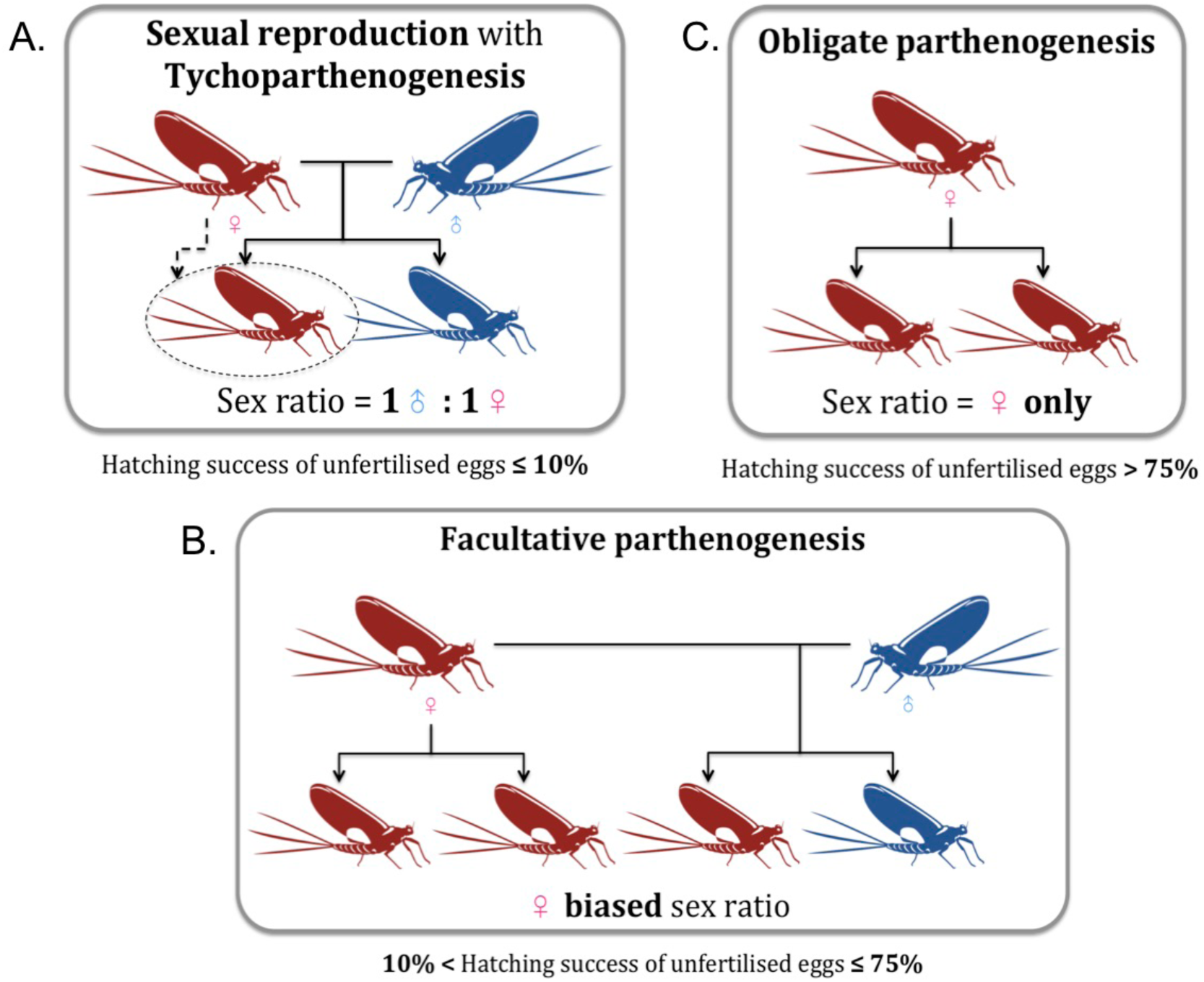
Three forms of female-producing (thelytokous) parthenogenesis in mayflies. A) Tychoparthenogenesis or “spontaneous parthenogenesis”, occurs in sexual species (typically less than 10% of unfertilised eggs develop through parthenogenesis). Given the low hatching success of unfertilised eggs, population sex ratios remain equal. B) Facultative parthenogenesis, when an egg may either be fertilised or develop parthenogenetically. The hatching success of unfertilised eggs in this case is intermediate (typically 10-75%), leading to female-biased population sex ratios. C) Under ‘obligate’ parthenogenesis, eggs always develop parthenogenetically and likely cannot be fertilised, with a hatching success typically higher than 75%. Only females are present in these populations. Note that an individual species can feature multiple forms of parthenogenesis, in the same or different populations.

## MATERIAL AND METHODS

### Data collection

The species list was compiled by collecting information from the literature on different websites: Google Scholar^1^, Web of Science^2^, Ephemeroptera Galactica^3^ and Ephemeroptera of the world^4^. Starting with four previous reviews (Degrange, 1960; Humpesch, 1980; Sweeney and Vannote, 1987; Funk *et al.*, 2010) that allowed us to compile a first list of 78 mayflies species studied for their reproductive mode, our survey combined with personal communications and observations generated a list of 136 species, as described in our database (Table 1, Appendix). When available for a given species, information on its geographical distribution, cytological mechanism of parthenogenesis and sex determination was included in the database (Appendix).

**Table 1.** Frequency of parthenogenesis and sex determination in Ephemeroptera taxa. Only families with at least one species studied for its parthenogenetic capacity are shown Seventeen families out of 42 (40.5%) have been studied. The the total numbers of described species have been adapted from Sartori and Brittain (2015) and updated (MS personal communication).

| Family | Parthenogenetic species | Sexual species | Total number of described species | % of parthenogenesis |  | Sex determination (2n+♀:♂). | References |
| --- | --- | --- | --- | --- | --- | --- | --- |
|  |  |  |  | among described species | among studied species |  |  |
| **Ameletidae** | 4 | 1 | 60 | 6.7 | 80.0 | 16+XX:XY | Morgan, 1911; Clemens, 1922; Needham, 1924; Traver, 1932; Katayama, 1939; Wolf, 1946; Burks, 1953; Sweeney and Vannote, 1982, 1987; Jazdzewska and Wojcieszek, 1997; Funk *et al.*, 2006; Reding, 2006 |
| **Baetidae** | 35 | 10 | 1076 | 3.3 | 77.8 | 8+XX:XY | Bengtsson, 1913; Dodds, 1923; McDunnough, 1925, 1936; Ide, 1930; Degrange, 1954, 1955, 1956, 1960; Tjønneland, 1960; Wolf, 1960; Hirvenoja, 1964; Uéno, 1966; Bohle, 1969; Gibbs, 1977; Kiauta and Mol, 1977; Bergman and Hilsenhoff, 1978; McCafferty and Morihara, 1979; Kluge, 1980; Harper and Harper, 1984; Sweeney and Vannote, 1984, 1987; Sivaramakrishnan *et al.*, 1991; Lowen and Flannagan, 1992; Sweeney *et al.*, 1993; Jackson and Sweeney, 1995; Harker, 1997; Salas and Dudgeon, 1999; Gattolliat and Sartori, 2000; Soldan and Putz, 2000; Funk *et al.*, 2006, 2008, 2010; Encalada and Peckarsky, 2007; Giberson *et al.*, 2007; Webb *et al.*, 2012; Martynov, 2013; ML pers. obs. |
| **Baetiscidae** | - | 1 | 12 | 0 | 0 | NA | Pescador, 1973; Pescador and Peters, 1974 |
| **Behningiidae** | - | 1 | 7 | 0 | 0 | NA | Peters and Peters, 1977; Harvey *et al.*, 1980; Sweeney and Vannote, 1982 |
| **Caenidae** | 5 | 4 | 258 | 1.9 | 55.6 | 6+XX:XO | Degrang, 1960; Froehlich, 1969; Mol, 1978; Gillies and Knowles, 1990; Da-Silva, 1993, 1998; Hofmann *et al.*, 1999; Soldan and Putz, 2000; Malzacher and Staniczek, 2007; FFS pers. com. |
| **Dipteromimidae** | - | 1 | 2 | 0 | 0 | NA | Takenaka *et al.*, 2019 |
| **Ephemerellidae** | 7 | 4 | 154 | 4.5 | 63.6 | 6, 14+XX:XY | Needham, 1924; Degrange, 1960; Kazlauskas, 1963; Fiance, 1978; Mol, 1978; Harper and Harper, 1982; Sweeney and Vannote, 1987; Soldan and Putz, 2000; Glazaczow, 2001; Funk *et al.*, 2008, 2010; Webb *et al.*, 2012 |
| **Ephemeridae** | 1 | 10 | 85 | 1.2 | 9.1 | 10, 12+XX:XO | Neave, 1932; Wolf, 1946; Hunt, 1951; Degrange, 1960; Britt, 1962; Thomforde and Fremling, 1968; Friesen and Flannagan, 1976; Mol, 1978; Sweeney and Vannote, 1987; Soldan and Putz, 2000; Funk *et al.*, 2010; Sekiné and Tojo, 2010a |
| **Heptageniidae** | 3 | 22 | 610 | 0.5 | 12 | 18+XX:XY | Degrang, 1955, 1960; Huff and McCafferty, 1974; McCafferty and Huff, 1974; Mingo, 1978; Mol, 1978; Humpesch, 1980; Salas and Dudgeon, 1999; Soldan and Putz, 2000; Ball, 2001, 2002; Funk *et al.*, 2010; ML pers. obs. |
| **Isonychiidae** | - | 1 | 35 | 0 | 0 | NA | Sweeney and Vannote, 1987 |
| **Leptophlebiidae** | 4 | 10 | 726 | 0.6 | 28.6 | 12-14+XX:XY | Degrang, 1960; Clifford *et al.*, 1979; Savage, 1986; Sweeney and Vannote, 1987; Jackson and Sweeney, 1995; Salas and Dudgeon, 1999; Soldan and Putz, 2000; Funk *et al.*, 2010 |
| **Oligoneuriidae** | - | 1 | 66 | 0 | 0 | 14+XX:XY | Degrang, 1960; Soldan and Putz, 2000 |
| **Palingeniidae** | 1 | - | 33 | 3 | 100 | NA | Landolt *et al.*, 1997 |
| **Polymitarcyidae** | 2 | 2 | 104 | 1.9 | 50 | 10, 14+XX:XO | Degrang, 1960; Britt, 1962; Tojo *et al.*, 2006; Sekiné and Tojo, 2010b, 2010a; Sekiné *et al.*, 2013, 2015 |
| **Potamanthidae** | - | 1 | 25 | 0 | 0 | 6+XX:XO | Degrang, 1960; Soldan and Putz, 2000 |
| **Prosopistomatidae** | 1 | 1 | 29 | 3.4 | 50 | NA | Degrang, 1960; Campbell and Hubbard, 1998 |
| **Siphonuridae** | 2 | 1 | 48 | 4.2 | 66.7 | 16+XX:XY | Degrang, 1954, 1955, 1960; Gibbs and Siebenmann, 1996; Soldan and Putz, 2000 |
| **Total** | **65** | **71\*** | **3666** | **1.8%** | **47.8%** |  |  |

\* Among the 71 sexual species, 38 can reproduce by tytoparthenogenesis (53.5%).

### Classification of parthenogenesis

To classify mayfly species by forms of parthenogenesis (Box 1), we focused on the hatching rate of unfertilised eggs. We defined two main categories according to this rate: “sexual species” (including species with tychoparthenogenesis), when less than 10% of unfertilised eggs hatch (typically 0-5.3% for population means), and “parthenogenetic species”, when hatching success of unfertilised eggs is higher than 10% (typically 18.5-97.3% for population means). Parthenogenetic species include all species with facultative parthenogenesis, ‘obligate’ parthenogenesis and mixed reproduction. Note that species with mixed reproduction have low population-average hatching success of unfertilised eggs when sexual and parthenogenetic females occur in sympatry (see results). We also considered species as parthenogenetic if female-only populations were reported in the literature, even if these species were not directly tested for their parthenogenetic capacity (this was the case for 18 of the 136 species in the database).

Within parthenogenetic species, we further distinguished ‘obligate’ from facultative parthenogens. Excluding rare events of sex in putatively obligate pathenogens is difficult (reviewed in Schurko *et al.*, 2009). We used the term obligate parthenogens for species where no males are known, or where rare males (typically <0.1% of all individuals) are most likely vestiges of sexual reproduction (van der Kooi and Schwander, 2014), indicating that parthenogenesis is the main form of reproduction. We also found very rare mentioning of deuterotoky (where both males and females are produced parthenogenetically). Specifically, the baetid species *Centroptilum luteolum*, *Acerpenna pygmaea*, *Acerpenna macdunnoughi* and *Anafroptilum semirufum* were inferred to be deuterotokous in breeding studies (Degrange, 1956; Funk *et al.*, 2010) because parthenogenetic broods contained high frequencies (2-17%) of males. Occasional deuterotoky is also the most likely explanation for the occurrence of rare males in ‘obligately’ parthenogenetic species as mentioned above. It is currently unclear how such males are produced (e.g., via environmental influences on sex determination or X-chromosome losses in species with XX/X0 sex determination). The few known deuterotokous species are counted as parthenogenetic species in our classification.

### Statistical analyses

We first verified that our classification into sexual (with or without tychoparthenogenesis) and parthenogenetic species (facultative, mixed and obligate) was biologically meaningful, by comparing the distribution of hatching successes of unfertilised eggs for different groups (Fig. 1A). We then tested whether species with high egg-hatching successes were often characterised by female-biased population sex ratios, by using a quasibinomial Generalised Linear Mixed Model (GLMM) with the R v.3.3.3 (R Development Core Team, 2017) ‘MASS’ (Venables and Ripley, 2002) and ‘car’ (Fox and Weisberg, 2011) packages. We then compared the prevalence of parthenogenetic species among mayfly families, using the most recent phylogeny available (Ogden *et al.*, 2019). Lack of phylogenetic information at lower taxonomic levels precluded further analyses. In order to obtain the most representative frequency estimate for the group, we also used the inventory of sexual species in our analyses. Indeed, the two available previous estimates of the frequency of parthenogenesis among animals (White, 1973; Vrijenhoek, 1998; van der Kooi *et al.*, 2017) assumed that all described species without evidence for parthenogenesis were sexual. However, this assumption severely underestimates the frequency of parthenogenesis. To account for this underestimation, we generated two frequency estimates, one using the total number of described mayfly species, and one using only species where the reproductive mode was studied. We thus tested whether variations in the frequency of parthenogenesis among families were explained by their phylogenetic relatedness by using binomial Generalised Linear Models (GLMs) and Tukey tests with the ‘multcomp’ (Hothorn *et al.*, 2008) package. In addition, we tested whether the frequency of parthenogenesis in mayflies varies among the six broad geographical regions: Nearctic, Palearctic, Neotropical, Afrotropical, Oriental and Australasian (see map in Table 2). Finally, we tested for potential trade-offs between parthenogenetic and sexual reproduction by studying hatching rate of fertilised and unfertilised eggs at the population level of a given species (Figure 5).

**Table 2.** World distribution of parthenogenesis in Ephemeroptera.

| Geographical region | Parthenogenetic species | Sexual species | Total number of described species per region | % of parthenogenesis among described species (left) among studied species (right) |  |
| --- | --- | --- | --- | --- | --- |
| NEA | 26 | 26 | 611 | 4.3 | 50.0 |
| PAL (west + east) | 23 | 44 | 839 | 2.7 | 34.3 |
| AFR | 5 | 2 | 642 | 0.8 | 71.4 |
| NEO | 7 | 0 | 902 | 0.8 | 100 |
| ORI | 3 | 0 | 440 | 0.7 | 100 |
| AUS + PAC | 2 | 0 | 299 | 0.7 | 100 |

NEA= Nearctic, NEO=Neotropical, PAL=Palearctic, AFR=Afrotropical, AUS=Australasian, PAC=Pacific islands. Adapted from Barber-James *et al.* (2008) and updated (MS pers. com.).

**Figure 1.**
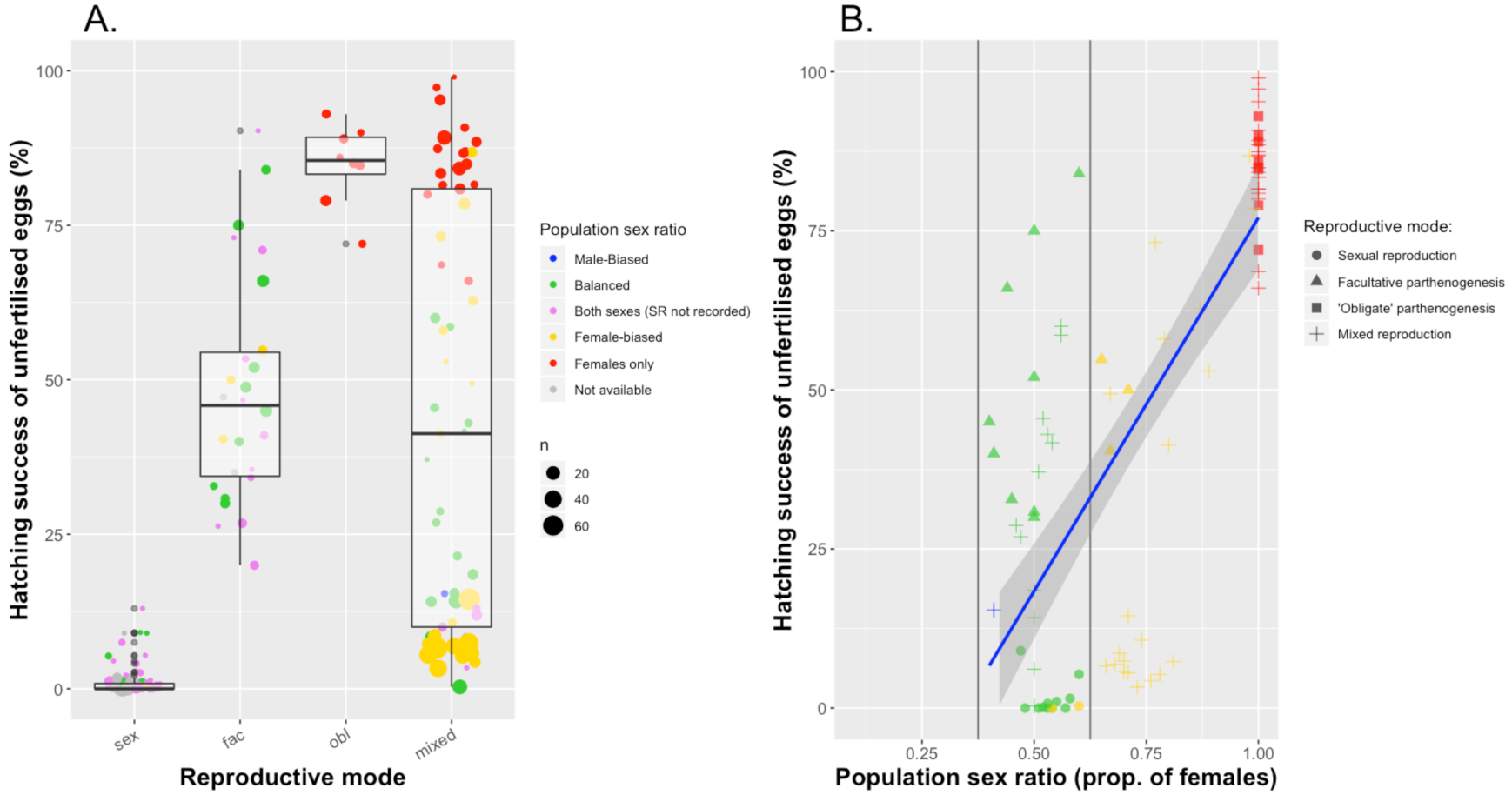
A) Population-average hatching successes of unfertilised eggs according to the species’ reproductive modes. sex: Sexual Reproduction, fac: Facultative parthenogenesis, obl: ‘Obligate’ parthenogenesis, mixed: Mixed reproduction (in sympatry and/or in allopatry). n: number of females tested for a given population. **B) Population-level correlation between unfertilised egg-hatching successes and sex ratios** (*Cor=0.72, p-value <0.001*).

## RESULTS

### Unfertilised egg-hatching successes and population sex ratios

Analysing the information we collected in our database revealed that the parthenogenetic capacity of females varied widely between and within populations (see Appendix for details). Nevertheless, our classification into sexual (with or without tychoparthenogenesis) and parthenogenetic species (facultative or obligate) is biologically meaningful, given the largely non-overlapping values for population-average hatching successes of unfertilised eggs (Figure 1). Note that some species with mixed reproduction show a low population-average hatching success of unfertilised eggs when sexual and parthenogenetic females occur in sympatry (e.g., average of 5.7% for one population of *Stenonema femoratum*, with egg-hatching successes varying among females from 0 to 77.9%). In order to determine whether a higher capacity for parthenogenesis translates into female-biased population sex ratios, we used species where both sex ratios and unfertilised egg-hatching successes were studied in the same populations. In these species, the parthenogenetic capacity and population sex ratios were significantly positively correlated (Fig. 1B, *cor=0.72, p-value <0.001*). The parthenogenetic capacity of females in strongly biased populations (>60% of females) was always very high (median: 83.4%, range: 40.4-97.3%), except for the species with mixed reproduction in sympatry as mentioned above (median: 7.8%, range: 3.3-15.5%). Conversely, unbiased population sex ratios were not indicative of species with obligate sexual reproduction, as they frequently comprised females with a high parthenogenetic capacity (Figure 1).

### Frequency of parthenogenesis among mayflies

Parthenogenesis occurs in all well-studied mayfly families (Table 1, Fig. 3). We were able to classify the reproductive mode of 136 species from 17 families (Table 1, see Appendix for details). Seventy-one of these species are sexual (from 16 families), and 38 of these are able to perform tychoparthenogenesis, while 65 species are parthenogenetic (from 11 families). Assuming the 3’666 described mayfly species without information concerning their reproductive mode are sexual, 1.8% of all mayfly species are able to reproduce parthenogenetically, which is at least an order of magnitude higher than the available estimate for vertebrates (0.1%, White, 1973; Vrijenhoek, 1998), and comparable to the frequencies in other arthropod orders. For example, the frequency of parthenogenesis in orders with haplodiploid sex determination varies from 0 to 1.5% (van der Kooi *et al.*, 2017). However, if one uses the frequency estimates based on the number of mayfly species studied for their reproductive mode (n=136), the estimated frequency of parthenogenesis reaches 47.8% (Fig. 2), being about 25 times higher. These findings suggest that half of the mayfly species might be able to reproduce parthenogenetically, or even, that most mayflies are able to reproduce at least by tychoparthenogenesis (75.7% of the studied species).

**Figure 2.**
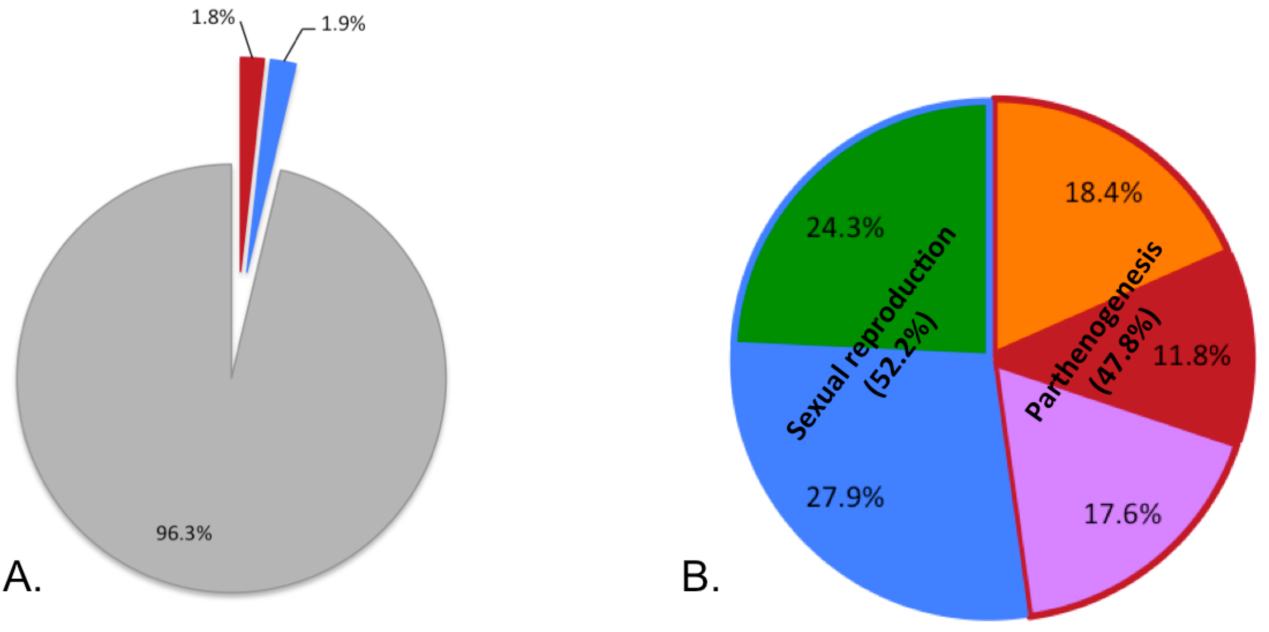
Frequency of parthenogenesis among described mayfly species (A) and among species studied for their reproductive mode (B). Less than four percent of the mayfly species have been studied for their reproductive mode. Parthenogenesis (warm colours): facultative (orange), ‘obligate’ (red) and mixed reproduction (purple). Sexual reproduction (cold colours): ‘obligate’ sexual reproduction (green) and sexual reproduction with tychoparthenogenesis (blue). At least 47.8% of the studied species are able to reproduce parthenogenetically.

**Figure 3.**
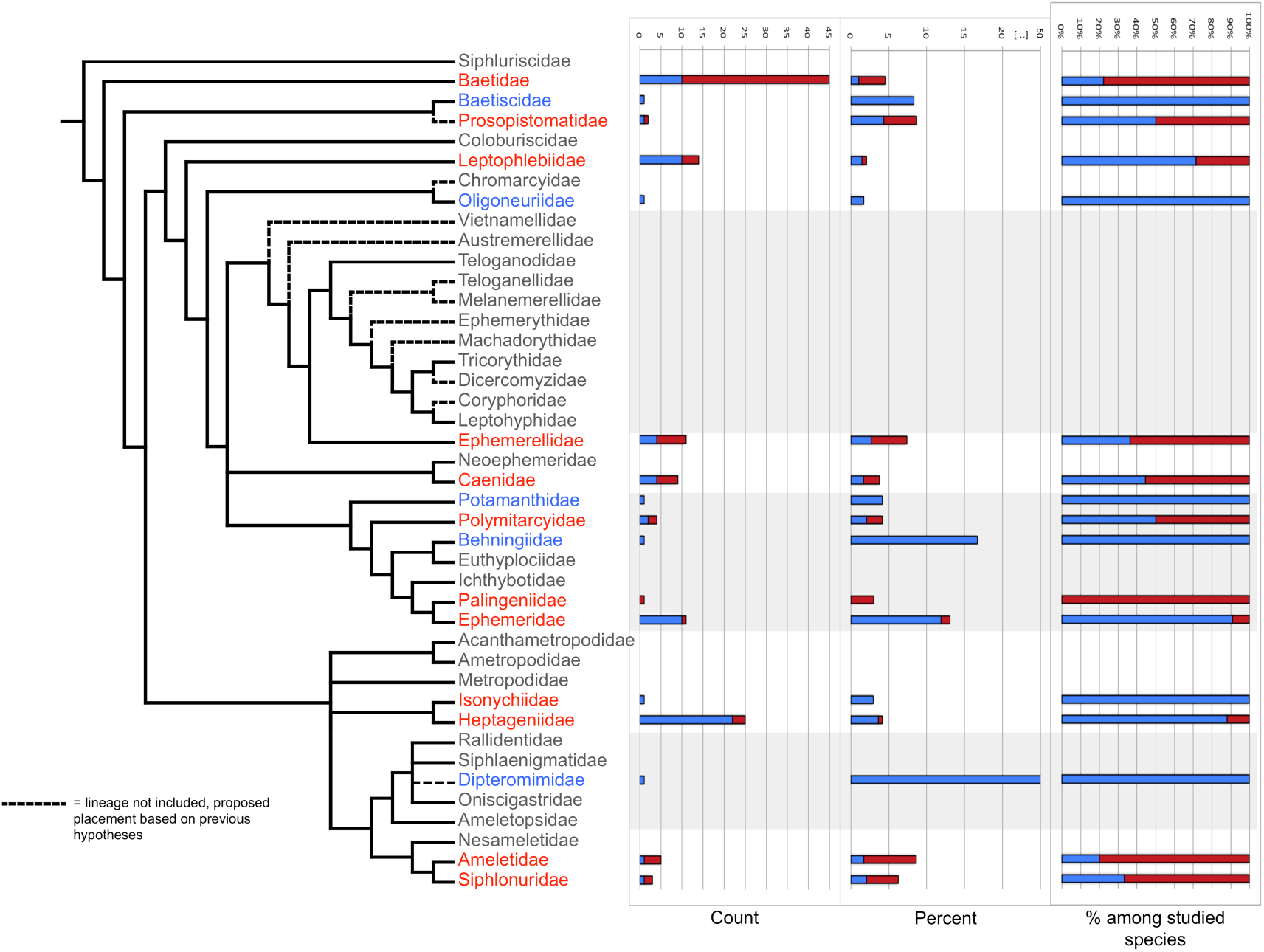
Phylogenetic distribution of parthenogenesis among mayfly families. Parthenogenetic species (facultative, obligate and mixed), Sexual species (with or whithout tychoparthenogenesis), Families without information regarding species’ reproduction. The families Heptageniidae, Leptophlebiidae and Ephemeridae show a low propensity for parthenogenesis, whereas the families Baetidae, Ameletidae and Ephemerellidae show a high propensity for parthenogenesis (see main text for details). Phylogeny adapted from Ogden *et al.* (2009, 2019).

Parthenogenesis occurs in an array of families and genera without any evidence for phylogenetic clustering (Fig. 3). A similar pattern is observed in haplodiploid taxa (van der Kooi *et al.*, 2017). These findings suggest that putative predispositions for the evolution of parthenogenesis do not have a deep phylogenetic inertia within the studied groups. However, clustering occurs at lower taxonomic levels as the proportion of parthenogenetic species varies significantly among mayfly families (*p-value <0.001*). Indeed, parthenogenetic mayfly species are rarer in the families Heptageniidae, Leptophlebiidae and Ephemeridae (<1.5% among the described mayfly species, or <30% among the one studied for their reproductive mode), than in the families Baetidae, Ameletidae and Ephemerellidae (>3.0% or >60%).

### Geographical parthenogenesis

The term geographical parthenogenesis is used when parthenogenetic populations are found at higher altitudes or latitudes than their sexual counterparts, in harsher environmental conditions or have wider distributions and/or ecological niches (Vandel, 1928). Such different distributions of sexual and asexual species could provide insights into ecological conditions that favour sex or parthenogenesis in natural populations, but quantitative comparisons are scarce (reviewed in Tilquin and Kokko, 2016).

Considering very broad geographical scales and among-species comparisons, we found no evidence for geographic clustering of parthenogenetic species. The six regions compared comprised approximately equal proportions of studied parthenogenetic species (Table 2, see Appendix for details). However, even if such differences existed, we would not be able to uncover them with the currently available data. Indeed, among the 136 studied species, 117 come from Nearctic and Palearctic regions (86%), with very little data available for the remaining regions of the world.

Considering within-species comparisons at smaller geographical scales, there are at least 18 mayfly species (27.7% of the parthenogenetic species) with both sexual and parthenogenetic populations (see Appendix for details). Two of them feature geographical parthenogenesis, but with distinct distribution differences between parthenogens and sexuals. Parthenogenetic populations of *Eurylophella funeralis* (Ephemerellidae) mostly occur at the periphery of the species ranges in North America (Sweeney and Vannote, 1987), while parthenogenetic populations of *Ephemerella notata* (Ephemerellidae) occur at lower latitudes than sexual ones in Poland (Glazaczow, 2001). One of the 18 species does not feature geographical parthenogenesis. Indeed, no geographical pattern is observed for *Ephoron shigae* (Polymitarcyidae), a species where sexual and parthenogenetic populations broadly overlap in Japan (Watanabe and Ishiwata, 1997). Finally, for 15 of the 18 species where sexual and parthenogenetic populations are known to occur in separate geographical areas, the number of described populations is too small to distinguish between a systematic distribution difference from a patchy distribution of sexual and parthenogenetic populations (Appendix).

More than two cases of geographical parthenogenesis likely exist in mayflies but are not detected because of a lack of studies, especially in the southern hemisphere. However, it seems that geographical parthenogenesis is not necessarily associated with particular ecological factors, as is the case in most other taxonomic groups studied thus far (Tilquin and Kokko, 2016).

### Cytological mechanisms of parthenogenesis in mayflies

In animals, different cytological mechanisms can underlie thelytokous parthenogenesis, which vary with respect to their consequences on heterozygosity in offspring (reviewed in Suomalainen *et al.*, 1987). In mayflies, some of these mechanisms have been identified or suggested, but studies remain scarce. Nevertheless, the available information suggests that obligate parthenogens use cytological mechanisms that potentially allow for maintenance of heterozygosity across generation (but see Jaron *et al.*, 2018), while facultative parthenogens are invariably automictic and produce parthenogenetic offspring that are highly homozygous relative to sexual offspring. Specifically, nine out of the 10 studied ‘obligately’ parthenogenetic mayflies are functionally clonal without a detected loss of heterozygosity between generations (Sweeney and Vannote, 1987; Sweeney *et al.*, 1993; Funk *et al.*, 2006, 2008, 2010). Three mechanisms can be responsible of the complete maintenance of heterozygosity between generations: apomixis (no meiosis occurs – mitotic parthenogenesis), endoduplication, or automixis with central fusion (without recombination). Which one(s) of these mechanisms occur in mayflies is currently unknown. The cytological mechanism of the remaining species, *E. shigae* in Japan, was suggested to be automixis with terminal fusion (Sekiné and Tojo, 2010b), indicating that some ‘obligate’ parthenogens might not be clonal.

All seven studied facultatively parthenogenetic mayflies are automictic (Appendix). Indeed, for the populations with both males and females of seven Baetidae species (*Acerpenna macdunnoughi*, *A. pygmaea*, *Anafroptlilum semirufum*, *Labiobaetis frondalis*, *Neocloeon alamance*, *Procloeon fragile*, *P. rivulare*) parthenogenesis appears to be automictic with terminal or central fusion (with recombination), given the partial loss of heterozygosity between generations, but further details are not known (Funk *et al.*, 2010). Finally, the cytological mechanism in *Ephoron eophilum* (Polymitarcyidae), a mostly sexual species with some facultatively parthenogenetic females (i.e., mixed reproduction in sympatry) is either automixis with terminal fusion (without recombination) or gamete duplication, where complete homozygosity is achieved in one generation (Sekiné *et al.*, 2015). Cytological mechanisms of parthenogenesis in mayflies clearly require additional studies, but the major mechanisms identified to date are summarised in Figure 4.

**Figure 4.**
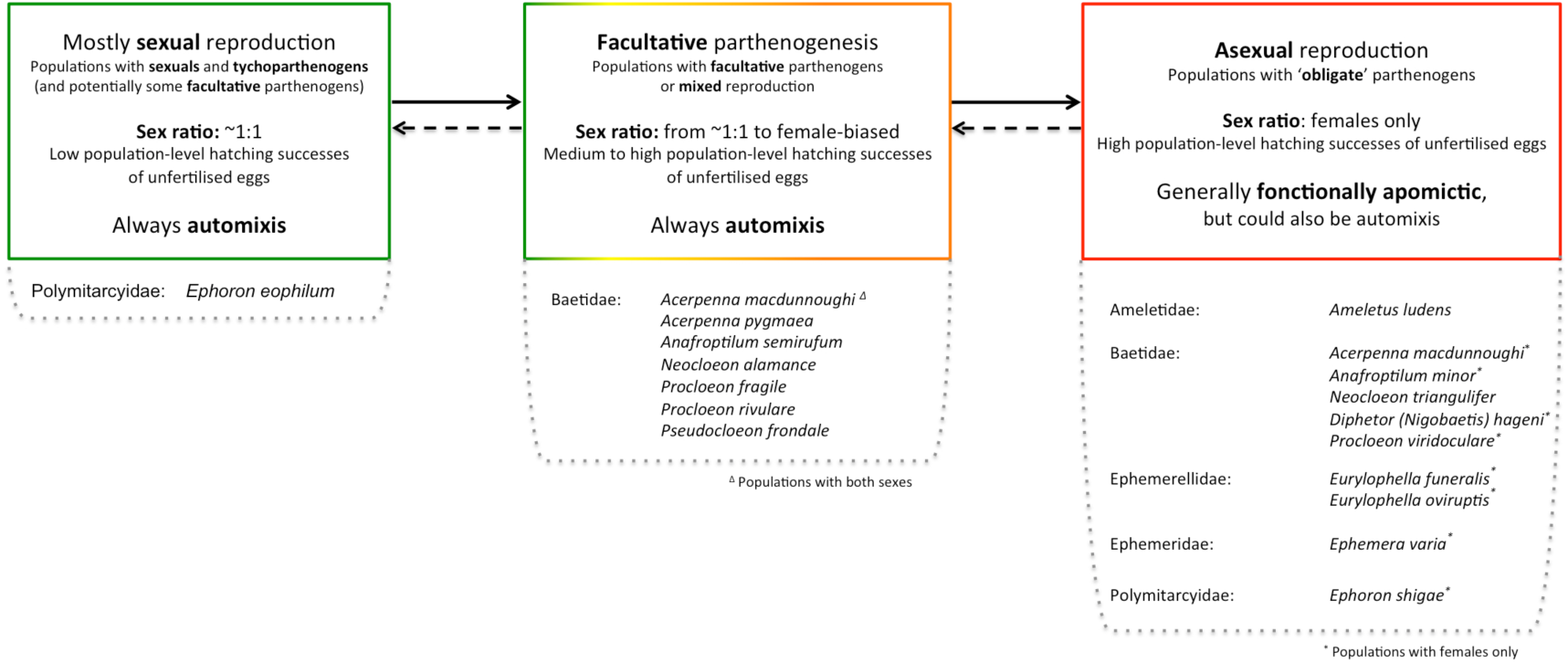
Cytological mechanisms identified to date in mayflies and possible transition between parthenogenesis forms.

Overall, mayfly species are better at reproducing sexually than asexually (measured as egg-hatching success, Fig. 5A, *p-value <0.001*). Only in obligate parthenogens is egg-hatching success decreased upon mating, presumably because (even partial) fertilisation interferes with normal development of asexual eggs. Furthermore, there is a significant negative correlation between hatching rate of fertilised and unfertilised eggs at the population level of a given species (Fig. 5B, *cor=-0.50, p-value=0.02*). This negative correlation suggests that there are trade-offs between parthenogenetic and sexual reproduction, meaning that improving the capacity for parthenogenesis may come at the cost of being less able to reproduce sexually, even in facultative parthenogens. If such a trade-off indeed exists, it could help explain why facultative parthenogenesis is extremely rare among animals in spite of its potential to combine the benefits of sexual and parthenogetetic reproduction.

**Figure 5.**
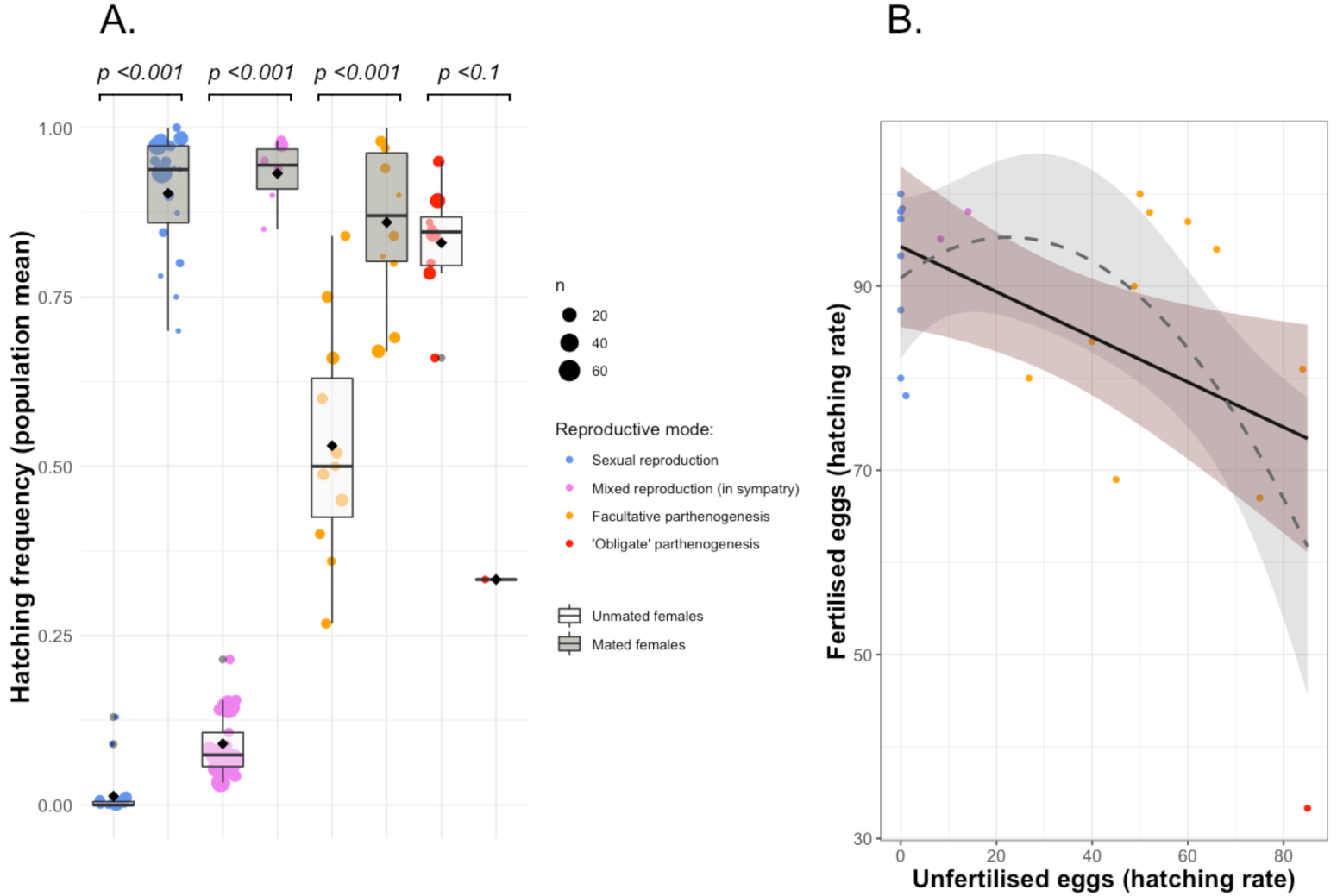
A) Hatching success of fertilised and unfertilised eggs for species with different reproductive modes and mating status. n: number of females tested for a given population; **B) Trade-off between parthenogenesis and sexual reproduction** of a given population (*cor=-0.50, p-value=0.02*).

### Origins of ‘obligate’ parthenogenesis in mayflies

There are at least four ways in which parthenogenetic lineages could arise from sexual species in animals: (1) Hybridisation between two sexual species, which is the major route to parthenogenesis in vertebrates (Avise *et al.*, 1992); (2) Contagious origin from pre-existing parthenogenetic lineages, where males produced by parthenogenetic species generate new lineages by mating with sexual females (e.g., in aphids Jaquiéry *et al.*, 2014; and water fleas Xu *et al.*, 2015); (3) Infection by microorganisms (e.g., *Wolbachia*, *Cardinium*, *Rickettsia*), mostly in species with haplodiploid sex determination (reviewed in Ma and Schwander, 2017) and (4) Spontaneous transitions through mutations, for example with tychoparthenogenesis as a first step (Carson *et al.*, 1957; Kramer and Templeton, 2001; Schwander and Crespi, 2009; Schwander *et al.*, 2010).

In mayflies, there is no evidence of parthenogenesis induced by hybridisation or endosymbiont infection, but there is very little data informing on the origins of parthenogenesis. An hybrid origin seems unlikely because it usually results in high levels of heterozygosity (recently reviewed in Jaron *et al.*, 2018) which is not the case for unisexual populations of the mayfly species studied so far (Sweeney and Vannote, 1987; Funk *et al.*, 2006; Sekiné and Tojo, 2010b). Endosymbiont induced parthenogenesis is also unlikely in mayflies because parthenogenesis in this group is often facultative, while endosymbiont infection normally causes obligate parthenogenesis (reviewed in Ma and Schwander, 2017). In addition, sex determination is male heterogamety (not haplodiploïdy, Table 1, see Appendix for details), further reducing the probability for endosymbiont-induced parthenogenesis.

In mayflies, it has been speculated that facultative and obligate parthenogenesis originates from tychoparthenogenesis (Sweeney and Vannote, 1987; Tojo *et al.*, 2006). Although this is a plausible hypothesis given how widespread tychoparthenogenesis is among mayflies (27.9% of the studied species, Fig. 2, see Appendix for details), there is no actual evidence for this suggestion. Indeed, there is currently very little information available that allows inferring how (facultative or obligate) parthenogenesis evolved in any of the known mayfly species. Nevertheless, because of their low dispersal ability and their short and fragile adult life, mayflies have restricted opportunities for reproduction, which may frequently generate situations of mate limitation in females. Mate limitation has been shown to favour parthenogenesis in other insect species (Schwander *et al.*, 2010), and is very likely to also select for parthenogenesis in mayflies, in spite of the probable trade-off with sexual reproduction we highlighted above.

### Fate of sexual traits in ‘obligate’ parthenogenetic mayflies

Sexual traits in asexual species decay more or less rapidly depending on whether they become costly or neutral upon transitions to parthenogenesis (reviewed in van der Kooi and Schwander, 2014). Selection favours the reduction of costly traits, contrary to neutral traits which decay via drift. For example, sexual traits which could decay in parthenogenetic females mayflies are: (1) pheromones, (2) the capacity to produce males, (3) the ability to fertilise their eggs, and (4) the synchrony of emergences.

Sex pheromones are chemical signals involved in mate choice (reviewed in Johansson and Jones, 2007) that can disappear in asexual lineages (e.g., Schwander *et al.*, 2013; Tabata Jun *et al.*, 2017). However, there are apparently no volatile pheromones in mayflies, with mate choice and species recognition based on visual signals and tactile recognition (Landolt *et al.*, 1997; MS pers. com.).

Occasional production of males has been reported in a range of ‘obligately’ parthenogenetic mayfly species (e.g., in *Ameletus ludens* Clemens, 1922; Needham, 1924; and in *Neocloeon triangulifer* Funk *et al.*, 2006, 2010) similar to most parthenogenetic species in other animal groups (van der Kooi and Schwander, 2014). In species with male heterogamety, male development in parthenogenetic lineages likely follows the accidental loss of an X chromosome during oogenesis (Schwander *et al.*, 2013). Accidental males produced by parthenogenetic females are often still able to fertilise eggs of females from sexual populations (van der Kooi and Schwander, 2014), but there is currently little information on the fertility of accidental males in mayflies. If fertile, as shown for two baetid species (Funk *et al.*, 2010), such males could potentially generates new lineages by matings with sexual females (i.e., contagious parthenogenesis as explained above), which could help explain the high frequency of parthenogenesis in mayflies.

The ability of parthenogenetic females to fertilise their eggs is unknown in mayflies overall, as only one species, the baetid *Neocloeon triangulifer*, has been studied thus far (Funk *et al.*, 2006). In this species, the ability to fertilise eggs is maintained at least at low levels. Viable progeny could be obtained by crossing *Neocloeon alamance* males (a sexual species with XY male heterogamety) with parthenogenetic *N. triangulifer* females. In such crosses, 66.6% of offspring were normal, clonal *N. tringulifer* females with high fertility, suggesting they were produced parthenogenetically from unfertilised eggs. However, the remaining 33.3% were genetically intermediate between the two species (as indicated by allozyme genotypes), suggesting they were hybrids produced from fertilised eggs. Approximately half of this hybrid progeny were females, perhaps triploid, with low fertility, the second half consisted of sterile gynandromorphs (with both male and females morphological characteristics). These findings suggest that even ‘obligately’ parthenogenetic mayfly species still produce haploid eggs, which could explain why there is always a small proportion of unfertilised eggs that never hatch (typically 3-22%), although more data are clearly needed.

Depending on how synchronous the emergences of both sexes are, the temporal windows to find a mate can be affected. Accordingly, obligate parthenogenesis might lead to a decay of the flight activity patterns in mayfly species. Tropical species of Trichoptera and Ephemeroptera appear to follow this theory (Tjønneland, 1970). However, this does not hold for several other mayfly species from temperate regions (e.g., Ameletidae: *Ameletus ludens*; Baetidae: *Neocloeon triangulifer*; Ephemerellidae: *Ephemerella notata* and *Eurylophella funeralis*), where emergence patterns of parthenogenetic mayflies are at least as synchronous as for sexual species (Sweeney and Vannote, 1982; Glazaczow, 2001). These findings might indicate that synchronisation of emergence is not costly in temperate regions, or that there are other factors such as predation that select for the maintenance of emergence synchrony (Sweeney and Vannote, 1982).

## CONCLUSION

We found evidence for parthenogenesis in at least 65 mayfly species, which represent as much as 1.8% of the 3’666 described species. However, this frequency is likely underestimated given that among the 136 species whose reproductive mode was studied, this value reaches 47.8%, currently the highest estimate known in non-cyclical parthenogenetic organisms. Parthenogenesis in mayflies thus appears to be widespread and is certainly an order of magnitude more frequent than in animal groups surveyed thus far. Among the 71 mayflies species found to reproduce sexually, 38 (53.5%) can produce offspring by accidental parthenogenesis (i.e., tychoparthenogenesis). Such accidental parthenogenesis could function as a pre-adaptation for facultative parthenogenesis, which may often be selected in mayflies because their short adult life frequently generates situations of mate limitation.

Additional studies focused on species from areas other than North America and Europe would be necessary to obtain a fully representative overview of the frequency of parthenogenesis in mayflies and for uncovering potential lineage-level or geographical-ecological correlates of parthenogenesis in this group. Additional studies are also needed regarding the cytological mechanisms and the origin of parthenogenesis in mayflies. In spite of these constraints, mayflies are currently clearly underappreciated for their value as outstanding model systems for testing benefits of sex in natural populations.

## Acknowledgement

We would like to thank Luca Sciuchetti for his contribution to review the literature and help for building the database. This study was supported by grants PP00P3_139013 and PP00P3_170627 from the Swiss NSF.

## Footnotes

1 https://scholar.google.com

2 https://apps.webofknowledge.com

3 www.ephemeroptera-galactica.com

4 www.insecta.bio.spbu.ru/z/Eph-spp/Contents.htm

## References

Agrawal AF (2006). Evolution of sex: Why do organisms shuffle their genotypes? Curr Biol 16: R696–R704.

Avise JC, Quattro JM, Vrijenhoek RC (1992). Molecular clones within organismal clones. In: Hecht MK, Wallace B, Macintyre RJ (eds) Evolutionary Biology, Springer: Boston, MA. Vol 26, pp 225–246.

Ball SL (2001). Tychoparthenogenesis and mixed mating in natural populations of the mayfly *Stenonema femoratum*. Heredity 87: 373–380.

Ball SL (2002). Population variation and ecological correlates of tychoparthenogenesis in the mayfly, *Stenonema femoratum*. Biol J Linn Soc 75: 101–123.

Barber-James HM, Gattolliat J-L, Sartori M, Hubbard MD (2008). Global diversity of mayflies (Ephemeroptera: Insecta) in freshwater. Hydrobiologia 595: 339–350.

Bauernfeind E, Moog O (2000). Mayflies (Insecta: Ephemeroptera) and the assessment of ecological integrity: a methodological approach. Hydrobiologia 422: 71–83.

Bengtsson S (1913). Undersökningar öfver äggen hos Ephemeriderna. Entomol Tidskr 34: 271–320 (+pl. 1-3).

Bergman E, Hilsenhoff W (1978). Parthenogenesis in mayfly genus *Baetis* (Ephemeroptera: Baetidae). Ann Entomol Soc Am 71: 167–168.

Bohle VHW (1969). Untersuchungen über die Embryonalentwicklung und die embryonale Diapause bei *Baetis vernus* (Curtis) und *Baetis rhodani* (Pictet) (Baetidae, Ephemeroptera). Zool Jahrb Abt Für Anat Ontog Tiere 86: 493–575.

Britt NW (1962). Biology of two species of Lake Erie mayflies, *Ephoron album* (Say) and *Ephemera simulans* Walker. (Ephemeroptera). Bull Ohio Biol Surv 1: 1–72.

Brittain JE, Sartori M (2009). Chapter 91 - Ephemeroptera: (Mayflies). In: Resh VH, Cardé RT (eds) Encyclopedia of Insects, Second Edition. Elsevier Academic Press: San Diego, CA, pp 328–334.

Burks BD (1953). The mayflies, or Ephemeroptera, of Illinois. Bull Ill Natl Hist Surv 26: 1–216.

Campbell IC, Hubbard MD (1998). A new species of *Prosopistoma* (Ephemeroptera: Prosopistomatidae) from Australia. Aquat Insects 20: 141–148.

Carson HL, Wheeler MR, Heed WB (1957). A parthenogenetic strain of *Drosophila mangabeirai* Malogolowkin. Genet Drosoph 5721: 115–122.

Clemens WA (1922). A parthenogenetic mayfly (*Ameletus ludens* Needham). Can Entomol 54: 77–78.

Clifford HF, Hamilton H, Killins BA (1979). Biology of the mayfly *Leptophlebia cupida* (Say) (Ephemeroptera: Leptophlebiidae). Can J Zool 57: 1026–1045.

Da-Silva ER (1993). Descrição do imago macho de *Caenis cuniana* Froehlich, com notas biológicas (Ephemeroptera, Caenidae). Rev Bras Zool 10: 413–416.

Da-Silva ER (1998). Estratégias de adaptação das espécies de Ephemeroptera às condições ambientais da Restinga de Maricá, Estado do Rio de Janeiro. In: Nessimian JL, Carvalho AL (eds) Ecologia de Insetos Aquáticos, Series Oecologia Brasiliensis, PPGE-UFRJ: Rio de Janeiro, Brasil. Vol 5, pp 29–40.

Degrange C (1954). Deux cas de parthénogenèse chez les Ephéméroptères - *Siphlonurus aestivalis* Eat. et *Centroptilum luteolum* Müll. Comptes Rendus Séances Académie Sci 239: 1082–1083.

Degrange C (1955). Nouveaux cas de parthénogenèse chez les Ephéméroptères. Comptes Rendus Séances Académie Sci 241: 1860–1861.

Degrange C (1956). La parthénogénèse facultative deutérotoque de *Centroptilum luteolum* (Müll.) (Ephéméroptères). Comptes Rendus Séances Académie Sci 243: 201–203.

Degrange C (1960). Recherches sur la reproduction des Ephéméroptères. Trav Lab Hydrobiol Piscic Univ Grenoble 50–51: 7–193.

Dodds GS (1923). Mayflies from Colorado. Descriptions of certain species and notes on others. Trans Am Entomol Soc 49: 93–114.

Edmunds GF Jr, McCafferty WP (1988). The mayfly subimago. Annu Rev Entomol 33: 509–529.

Encalada AC, Peckarsky BL (2007). A comparative study of the costs of alternative mayfly oviposition behaviors. Behav Ecol Sociobiol 61: 1437–1448.

Fiance S (1978). Effects of pH on the biology and distribution of *Ephemerella funeralis* (Ephemeroptera). Oikos 31: 332–339.

Fox J, Weisberg S (2011). An {R} Companion to Applied Regression, Second Edition. Sage: Thousand Oaks, CA.

Friesen MK, Flannagan JF (1976). Parthenogenesis in the burrowing mayfly *Hexagenia rigida* (Ephemeroptera). Can Entomol 108: 1295–1295.

Froehlich CG (1969). *Caenis cuniana* sp.n., a parthenogenetic mayfly. Beitr Zur Neotropischen Fauna 6: 103–108.

Funk DH, Jackson JK, Sweeney BW (2006). Taxonomy and genetics of the parthenogenetic mayfly *Centroptilum triangulifer* and its sexual sister *Centroptilum alamance* (Ephemeroptera: Baetidae). J North Am Benthol Soc 25: 417–429.

Funk DH, Jackson JK, Sweeney BW (2008). A new parthenogenetic mayfly (Ephemeroptera: Ephemerellidae: *Eurylophella* Tiensuu) oviposits by abdominal bursting in the subimago. J North Am Benthol Soc 27: 269–279.

Funk DH, Sweeney BW, Jackson JK (2010). Why stream mayflies can reproduce without males but remain bisexual: a case of lost genetic variation. J North Am Benthol Soc 29: 1258–1266.

Gattolliat J-L, Sartori M (2000). *Guloptiloides*: an extraordinary new carnivorous genus of Baetidae (Ephemeroptera). Aquat Insects 22: 148–159.

Gibbs KE (1977). Evidence for obligatory parthenogenesis and its possible effect on emergence period of *Cloeon triangulifer* (Ephemeroptera: Baetidae). Can Entomol 109: 337–340.

Gibbs KE, Siebenmann M (1996). Life history attributes of the rare mayfly *Siphlonisca aerodromia* Needham (Ephemeroptera: Siphlonuridae). J North Am Benthol Soc 15: 95–105.

Giberson DJ, Burian SK, Shouldice M (2007). Life history of the northern mayfly *Baetis bundyae* in Rankin Inlet, Nunavut, Canada, with updates to the list of mayflies of Nunavut. Can Entomol 139: 628–642.

Gillies MT, Knowles RJ (1990). Colonization of a parthenogenetic mayfly (Caenidae: Ephemeroptera) from Central Africa. In: Campbell IC (ed) Mayflies and Stoneflies: Life histories and biology, Series Entomologica. Springer: Dordrecht. Vol 44, pp 341–345.

Glazaczow A (2001). Parthenogenetic and bisexual populations of *Ephemerella notata* Eat. in Poland. In: Domínguez E (ed) Trends in Research in Ephemeroptera and Plecoptera, Springer: Boston, MA, pp 227–231.

Harker JE (1997). The role of parthenogenesis in the biology of two species of mayfly (Ephemeroptera). Freshw Biol 37: 287–297.

Harper PP, Harper F (1982). Mayfly communities in a Laurentian watershed (Insecta; Ephemeroptera). Can J Zool 60: 2828–2840.

Harper F, Harper PP (1984). Phenology and distribution of mayflies in a southern Ontario lowland stream. In: Landa V, Soldán T, Tonner M (eds) Proceedings of the Fourth International Conference on Ephemeroptera, Institute of Entomology, Czechoslovak Academy of Sciences, České Budějovice, pp 243–251.

Harvey RS, Vannote RL, Sweeney BW (1980). Life history, developmental processes, and energetics of the burrowing mayfly *Dolania americana*. In: Flannagan JF, Marshall KE (eds) Advances in Ephemeroptera Biology, Plenum Press: New York, NY, pp 211–230.

Hirvenoja M (1964). Studien über die Wasserinsekten in Riihimäki (Südfinnland). IV: Ephemeroptera, Odonata, Hemiptera, Lepidoptera und Coleoptera. Ann Entomol Fenn 30: 65–93.

Hofmann C, Sartori M, Thomas A (1999). Les Ephéméroptères (Ephemeroptera) de la Guadeloupe (petites Antilles françaises). Mém Société Vaudoise Sci Nat 20: 1–96.

Hothorn T, Bretz F, Westfall P (2008). Simultaneous Inference in General Parametric Models. Biom J 50: 346–363.

Huff BL, McCafferty WP (1974). Parthenogenesis and experimental reproductive biology in four species of the mayfly genus *Stenonema*. Wasmann J Biol 32: 247–254.

Humpesch UH (1980). Effect of temperature on the hatching time of parthenogenetic eggs of five *Ecdyonurus spp*. and two *Rhithrogena spp*. (Ephemeroptera) from Austrian streams and English rivers and lakes. J Anim Ecol 49: 927–937.

Hunt BP (1951). Reproduction of the burrowing mayfly, *Hexagenia limbata* (Serville), in Michigan. Fla Entomol 34: 59–70.

Ide FP (1930). Contribution to the biology of Ontario mayflies with descriptions of new species. Can Entomol 62: 204–213, 218-231.

Jackson JK, Sweeney BW (1995). Egg and larval development times for 35 species of tropical stream insects from Costa Rica. J North Am Benthol Soc 14: 115–130.

Jalvingh K, Bast J, Schwander T (2016). Evolution and Maintenance of Sex. In: Kliman RM (ed) Encyclopedia of Evolutionary Biology, Elsevier Academic Press: Oxford, UK, pp 89–97.

Jaquiéry J, Stoeckel S, Larose C, Nouhaud P, Rispe C, Mieuzet L, et al. (2014). Genetic control of contagious asexuality in the Pea Aphid. PLoS Genet 10: e1004838.

Jaron KS, Bast J, Ranallo-Benavidez TR, Robinson-Rechavi M, Schwander T (2018). Genomic features of asexual animals. bioRxiv: 497495.

Jazdzewska T, Wojcieszek A (1997). *Metreletus balcanicus* (Ulmer, 1920) (Ephemeroptera) in Poland with notes on its ecology and biology. Pol Pismo Entomol 66: 9–16.

Johansson BG, Jones TM (2007). The role of chemical communication in mate choice. Biol Rev 82: 265–289.

Katayama H (1939). The sex chromosomes of a may-fly, *Ameletus costalis* Mats. (Ephemerida). Jpn J Genet 15: 139–144.

Kazlauskas RS (1963). [New and little-known mayflies (Ephemeroptera) from the fauna of the USSR]. Entomol Obozr 42: 582–593 (in Russian).

Kiauta B, Mol AWM (1977). Behaviour of the spermatocyte chromosomes of the mayfly, *Cloeon dipterum* (Linnaeus, 1761) s.1. (Ephemeroptera: Baetidae) with a note on the cytology of the order. Genen Phaenen 19: 31–39.

Kluge NJ (1980). [To the knowledge of mayflies (Ephemeroptera) of Taimyr National District]. Entomol Obozr 59: 561–579 (in Russian).

Knopp M, Cormier R (1997). Mayflies: An angler’s study of trout water Ephemeroptera, First Edition. Greycliff Publishing Company: Helena, MT.

van der Kooi CJ, Matthey‐Doret C, Schwander T (2017). Evolution and comparative ecology of parthenogenesis in haplodiploid arthropods. Evol Lett 1: 304–316.

van der Kooi CJ, Schwander T (2014). On the fate of sexual traits under asexuality. Biol Rev 89: 805–819.

Kramer MG, Templeton AR (2001). Life-history changes that accompany the transition from sexual to parthenogenetic reproduction in *Drosophila mercatorum*. Evolution 55: 748–761.

Landolt P, Sartori M, Studemann D (1997). *Palingenia longicauda* (Ephemeroptera: Palingeniidae): From mating to the larvulae stage. In: Landolt P, Sartori M (eds) Ephemeroptera & Plecoptera: Biology-Ecology-Systematics, MTL: Fribourg, Switzerland, pp 15–20.

Lehtonen J, Jennions MD, Kokko H (2012). The many costs of sex. Trends Ecol Evol 27: 172–178.

Lowen RG, Flannagan JF (1992). Nymphs and imagoes of four North American species of *Procloeon* Bengtsson with description of a new species (Ephemeroptera, Baetidae). Can Entomol 124: 97–108.

Ma W-J, Schwander T (2017). Patterns and mechanisms in instances of endosymbiont-induced parthenogenesis. J Evol Biol 30: 868–888.

Malzacher P, Staniczek AH (2007). *Caenis vanuatensis*, a new species of mayflies (Ephemeroptera: Caenidae) from Vanuatu. Aquat Insects 29: 285–295.

Martynov AV (2013). The life cycles of mayflies of the eastern Ukraine. Subfamily *Baetinae* (Ephemeroptera: Baetidae). Vestn Zool 47: 35–44.

McCafferty WP, Huff BL (1974). Parthenogenesis in the mayfly *Stenonema fermoratum* (Say), Ephemeroptera: Heptageniidae. Entomol News 85: 76–80.

McCafferty WP, Morihara DK (1979). The male of *Baetis macdunnoughi* Ide and notes on parthenogenetic populations within *Baetis* (Ephemeroptera: Baetidae). Entomol News 90: 26–28.

McDunnough J (1925). The Ephemeroptera of Covey Hill, Que. Trans R Soc Can 19: 207–223.

McDunnough J (1936). A new Arctic baetid (Ephemeroptera). Can Entomol 68: 33–34.

Mingo TM (1978). Parthenogenesis in the mayfly *Stenacron interpunctatum frontale* (Burks) (Ephemeroptera: Heptageniidae). Entomol News 89: 46–50.

Mol AWM (1978). Notes on the chromosomes of some West European Ephemeroptera. Chromosome Inf Serv 24: 10–12.

Morgan AH (1911). May-flies of Fall Creek. Ann Entomol Soc Am 4: 93–119.

Neave F (1932). A study of the May flies (Hexagenia) of Lake Winnipeg. Contrib Can Biol Fish 7: 177–201.

Needham JG (1924). The male of the parthenogenetic May-fly, *Ameletus ludens*. Psyche (Stuttg) 31: 308–310.

Ogden TH, Breinholt JW, Bybee SM, Miller D, Sartori M, Shiozawa D, et al. (2019). Mayfly phylogenomics: Initial evaluation of anchored hybrid enrichment data for the order Ephemeroptera. Zoosymposia in press.

Ogden TH, Gattolliat J-L, Sartori M, Staniczek AH, Soldán T, Whiting MF (2009). Towards a new paradigm in mayfly phylogeny (Ephemeroptera): combined analysis of morphological and molecular data. Syst Entomol 34: 616–634.

Otto SP (2009). The evolutionary enigma of sex. Am Nat 174: S1–S14.

Pescador ML (1973). The ecology and life history of *Baetisca rogersi* Berner (Ephemeroptera: Baetiscidae). In: Peters WL, Peters JG (eds) Proceedings of the First International Conference on Ephemeroptera, Florida, A&M Univ., 17-20 August 1970. Leiden, Brill, pp 211–215.

Pescador ML, Peters WL (1974). The life history and ecology of *Baetisca rogersi* Berner (Ephemeroptera: Baetiscidae). Bull Fla State Mus Biol Sci 17: 151–209.

Peters WL, Peters JG (1977). Adult life and emergence of *Dolania americana* in northwestern Florida (Ephemeroptera: Behningiidae). Int Rev Gesamten Hydrobiol 62: 409–438.

R Development Core Team (2017). R: A language and environment for statistical computing. R foundation for Statistical Computing. Vienna, Austria.

Reding J-P (2006). Notes faunistiques sur *Metreletus balcanicus* (Insecta: Ephemeroptera) et *Ironoquia dubia* (Insecta: Trichoptera), deux espèces d’insectes aquatiques du Jura nouvelles pour la Suisse. Bull Société Neuchâtel Sci Nat 129: 73–86.

Ross L, Hardy NB, Okusu A, Normark BB (2013). Large population size predicts the distribution of asexuality in scale insects. Evolution 67: 196–206.

Salas M, Dudgeon D (1999). Parthenogenesis in some Hong Kong mayflies (Ephemeroptera). Mem Hong Kong Nat Hist Soc 22: 165–169.

Sartori M, Brittain JE (2015). Chapter 34 - Order Ephemeroptera. In: Thorp JH, Rogers DC (eds) Ecology and General Biology: Thorp and Covich’s Freshwater Invertebrates, Elsevier Academic Press: Boston, MA. Vol 1, pp 873–891.

Savage HM (1986). Systematics of the *Terpides* lineage from the Neotropics: Definition of the *Terpides* lineage, methods, and revision of *Fittkaulus* Savage & Peters. Spixiana 9: 255–270.

Schurko AM, Neiman M, Logsdon JM (2009). Signs of sex: what we know and how we know it. Trends Ecol Evol 24: 208–217.

Schwander T, Crespi BJ (2009). Multiple direct transitions from sexual reproduction to apomictic parthenogenesis in *Timema* stick insects. Evolution 63: 84–103.

Schwander T, Crespi BJ, Gries R, Gries G (2013). Neutral and selection-driven decay of sexual traits in asexual stick insects. Proc R Soc B Biol Sci 280: 20130823.

Schwander T, Vuilleumier S, Dubman J, Crespi BJ (2010). Positive feedback in the transition from sexual reproduction to parthenogenesis. Proc R Soc B Biol Sci 277: 1435–1442.

Sekiné K, Hayashi F, Tojo K (2013). Phylogeography of the East Asian polymitarcyid mayfly genus *Ephoron* (Ephemeroptera: Polymitarcyidae): a comparative analysis of molecular and ecological characteristics. Biol J Linn Soc 109: 181–202.

Sekiné K, Tojo K (2010a). Potential for parthenogenesis of virgin females in a bisexual population of the geographically parthenogenetic mayfly *Ephoron shigae* (Insecta: Ephemeroptera, Polymitarcyidae). Biol J Linn Soc 99: 326–334.

Sekiné K, Tojo K (2010b). Automictic parthenogenesis of a geographically parthenogenetic mayfly, *Ephoron shigae* (Insecta: Ephemeroptera, Polymitarcyidae). Biol J Linn Soc 99: 335–343.

Sekiné K, Tojo K, Bae YJ (2015). Facultative parthenogenesis in the burrowing mayfly, *Ephoron eophilum* (Ephemeroptera: Polymitarcyidae) with an extremely short alate stage. Eur J Entomol 112: 606–612.

Sivaramakrishnan KG, Sridhar S, Rajarajan PA (1991). Effect of temperature on hatching of parthenogenetic eggs of *Baetis geminatus* Müller-Liebenan & Hubbard, 1985 from south India (Ephemeroptera; Baetidae). Opusc Zool Flum 69: 1–8.

Soldan T, Putz M (2000). Karyotypes of some Central European mayflies (Ephemeroptera) and their contribution to phylogeny of the order. Acta Soc Zool Bohemicae 64: 437–445.

Suomalainen E, Saura A, Lokki J (1987). Cytology and evolution in parthenogenesis. CRC Press, Boca Raton: Florida.

Sweeney BW, Funk DH, Standley LJ (1993). Use of the stream mayfly *Cloeon triangulifer* as a bioassay organism: Life-history response and body burden following exposure to technical chlordane. Environ Toxicol Chem 12: 115–125.

Sweeney BW, Vannote RL (1982). Population synchrony in mayflies: A predator satiation hypothesis. Evolution 36: 810–821.

Sweeney BW, Vannote RL (1984). Influence of food quality and temperature on life-history characteristics of the parthenogenetic mayfly, *Cloeon triangulifer*. Freshw Biol 14: 621–630.

Sweeney BW, Vannote RL (1987). Geographic parthenogenesis in the stream mayfly *Eurylophella funeralis* in eastern North America. Holarct Ecol 10: 52–59.

Tabata Jun, Ichiki Ryoko T., Moromizato Chie, Mori Kenji (2017). Sex pheromone of a coccoid insect with sexual and asexual lineages: fate of an ancestrally essential sexual signal in parthenogenetic females. J R Soc Interface 14: 20170027.

Takenaka M, Sekiné K, Tojo K (2019). The first establishment of “hand-pairing” cross-breeding method for the most ancestral wing acquired insect group. Zoolog Sci 36: 136–140.

Thomforde LL, Fremling CR (1968). Synchronous emergence of *Hexagenia bilineata* mayflies in the laboratory. Ann Entomol Soc Am 61: 1235–1239.

Tilquin A, Kokko H (2016). What does the geography of parthenogenesis teach us about sex? Philos Trans R Soc B Biol Sci 371: 20150538.

Tjønneland A (1960). The flight activity of mayflies as expressed in some East African species. Årb Univ Bergen Mat-Naturv Ser 1: 1–88.

Tjønneland A (1970). A possible effect of obligatory parthenogenesis on the flight activity of some tropical larvo-aquatic insects. Årb Univ Bergen Mat-Naturv Ser 3: 1–7.

Tojo K, Sekiné K, Matsumoto A (2006). Reproductive mode of the geographic parthenogenetic mayfly *Ephoron shigae*, with findings from some new localities (Insecta: Ephemeroptera, Polymitarcyidae). Limnology 7: 31–39.

Traver JR (1932). Mayflies of North Carolina. J Elisha Mitchell Sci Soc 47: 85–161 (+pl. 5-12), 163-236.

Uéno M (1966). Mayflies (Ephemeroptera) collected by the Kyoto University Pamir-Hindukush Expedition 1960. Results Kyoto Univ Sci Exped Karakoram Hindukush 1955 Kyoto 8: 299–326.

Vandel A (1928). La parthénogenèse géographique. Contribution à l’étude biologique et cytologique de la parthénogenèse naturelle. Bull Biol Fr Belg 62: 164–281.

Venables WN, Ripley BD (2002). Modern Applied Stastistics with S, Fourth Edition. Springer: New York, NY.

Vrijenhoek RC (1998). Animal clones and diversity: Are natural clones generalists or specialists? BioScience 48: 617–628.

Watanabe NC, Ishiwata S-I (1997). Geographic distribution of the mayfly, *Ephoron shigae* in Japan, with evidence of geographic parthenogenesis (Insecta: Ephemeroptera: Polymitarcyidae). Jpn J Limnol 58: 15–25.

Webb JM, Jacobus LM, Funk DH, Zhou X, Kondratieff B, Geraci CJ, et al. (2012). A DNA barcode library for North American Ephemeroptera: Progress and prospects. PLoS ONE 7: e38063.

White MJD (1973). Animal cytology and evolution, Third Edition. Cambridge University Press: London, UK.

Wolf E (1946). Chromosomenuntersuchungen an insekten. Z Für Naturforschung 1: 108–109.

Wolf E (1960). Zur karyologie der eireifung und furchung bei *Cloeon dipterum* L. (Bengtsson) (Ephemerida, Baetidae). Biol Zentralblatt 79: 153–198.

Xu S, Spitze K, Ackerman MS, Ye Z, Bright L, Keith N, et al. (2015). Hybridization and the origin of contagious asexuality in *Daphnia pulex*. Mol Biol Evol 32: 3215–3225.

