## Appendix for "Extremely widespread parthenogenesis and a trade-off between alternative forms of reproduction in mayflies (Ephemeroptera)"

| Systematic | | | Geography | | | | Reproductive mode | | Population sex ratio | Fecundity | Unfertilised eggs | | | Fertilised eggs | | | | | | |
|---|---|---|---|---|---|---|---|---|---|---|---|---|---|---|---|---|---|---|---|---|
| Family | *Species name* | Synonym(s) used | General Distribution | Country or State (studied populations) | County or Region (studied populations) | GPS coordinates (lat, lon) | "Class" of parthenogenesis (Species level) | Type of parthenogenesis (Population level) | Field sex ratio(s): prop. of females | number of eggs per female - mean (range, n) | Hatching rate (%) of unfertilised eggs - mean (range, n) | Development time (days) of unfertilised eggs | Hatching length of unfertilised eggs | Hatching rate (%) of fertilised eggs - mean (range, n) | Development time (days) of fertilised eggs | Hatching length of fertilised eggs | Cytological mechanisms | Karyotype and Sex determination (2n+♀:♂) | Comments | Reference(s) |
| Ameletidae | *Ameletus cryptostimulus* | | NEA | Eastern North America | | | **Sexual** | tycho. | both sexes | | 0.1 (0-0.1) | | | | | | | *Ameletus costalis*: 16 + XX:XY (Katayama, | | Sweeney&Vannote (1987) |
| Ameletidae | *Ameletus lineatus* | | NEA | North Carolina; Illinois | | several populations | **"Obligate"** | obl. | females only | | | | | | | | | | | Traver (1932); Burks (1953) |
| Ameletidae | *Ameletus ludens* | | NEA | New York | Ithaca | Fall Creek | **"Obligate"** | obl. | females only | | hatching | 5 months (diapause) | | | | | | | only two males founc | Morgan (1911); Clemens (1922); Needham (1924) |
| Ameletidae | *Ameletus ludens* | | NEA | Pennsylvania | | White Clay Creek (W | **"Obligate"** | obl. | females only: 1.00 | 223-402 | 90 (89-91) | | | | | | | | many years, hundred | Sweeney&Vannote (1982); Funk et al. (2006) |
| Ameletidae | *Ameletus tertius* | *Ameletus sp. near tertius* | NEA | Eastern North America | | | **Facultative** | fac. | both sexes | | 20 (20-22) | | | | | | | | | Sweeney&Vannote (1987) |
| Ameletidae | *Metreletus balcanicus* | *Ameletus hessei; Metreletus* | West PAL | France; Czech Republic; Germany; Switzerland | | several populations | **Mixed (allopatry)** | mostly obl. | strongly female-biased | | | | | | | | | | gynandromorphs | Jazdzewska&Wojcieszek (1997); Reding (2006); Tomáš Soldán (personal communication) |
| Ameletidae | *Metreletus balcanicus* | | West PAL | Poland; Germany; Belgium, Hungary; Slovakia | | several populations | **Mixed (allopatry)** | sex.? | balanced | | | | | | | | | | | reviewed in Jazdzewska&Wojcieszek (1997) |
| Baetidae | *Acentrella sinaica* | *Baetis subatrebatinus* | West PAL | France | Haute-Savoie | Le Fier, au pont de N | **Facultative** | fac. | both sexes | 735 | 26.3 (n=1) | 21-52d at 15°C | | | | | | | | Degrange (1960) |
| Baetidae | *Acentrella turbida* | *Pseudocloeon carolina* | NEA | Quebec; Arkansas; Colorado | | several populations | **Mixed (allopatry)** | obl. | females only | | | | | | | | | | | McDunnough (1925); McCafferty&Provonsha (1978); Encalada&Peckarsky (2007) |
| Baetidae | *Acentrella turbida* | *Pseudocloeon turbidum* | NEA | Alberta; Manitoba; Maine; British Columbia; O | | several populations | **Mixed (allopatry)** | sex.? | both sexes | | | | | | up to 9 months | | | | | McDunnough (1924); Cobb (1991); Burian & Gibbs (1991); McCafferty et al. (1994); Baumgardner et al. (1997); Webb et al. (2012) |
| Baetidae | *Acerpenna macdunnoughi* | *Baetis macdunnoughi* | NEA | Wisconsin | Marquette Co. | Lawrence Creek | **Mixed** | obl. | females only: 1.00 | | 95.3 (87-100, n=10) | 8.8d at 19-22°C (8-10, n=9); 15d at 13-15°C (n=1) | | | | | | | thelytokous parthenc | Bergman&Hilsenhoff (1978) |
| Baetidae | *Acerpenna macdunnoughi* | *Baetis macdunnoughi* | NEA | Wisconsin | Adams Co. | Carter Creek | **Mixed** | obl. | females only: 1.00 | | 66 (57-98, n=4) | 8.8d at 19-21°C (9-10, n=4) | | | | | | | thelytokous parthenc | Bergman&Hilsenhoff (1978) |
| Baetidae | *Acerpenna macdunnoughi* | *Baetis macdunnoughi* | NEA | Ontario, Quebec | | several populations | **Mixed** | obl. | females only | | | | | | | | | | | Ide (1937); Harper&Harper (1982, 1984) |
| Baetidae | *Acerpenna macdunnoughi* | | NEA | Pennsylvania | Chester Co. | White Clay Creek (W | **Mixed** | obl. | females only | | | | | | | | clonal: suggested apomixis or automixis (without recom | | | Funk et al. (2006, 2010) |
| Baetidae | *Acerpenna macdunnoughi* | | NEA | New York | Sullivan Co. | Neversink River: 41.9 | **Mixed** | fac. | balanced: 0.56 (n=34) | | 60 (34-83, n=8) | about twice as long as fertilised at 20°C | | forced copulation: 97 (95-99, n=4) | | | suggested automixis with central f | | 4% of males produce | Funk et al. (2010) |
| Baetidae | *Acerpenna macdunnoughi* | *Baetis macdunnoughi* | NEA | Illinois, Indiana | south of this 2 states | | **Mixed** | sex.? | balanced | | | | | | | | | | | McCafferty&Morihara (1979) |
| Baetidae | *Acerpenna pygmaea* | | NEA | Alabama | Clay Co. | Cheaha Creek: 33.45 | **Facultative** | fac. | balanced: 0.6 (n=15) | | 84 (72-95, n=6) | about twice as long as fertilised at 20°C | | forced copulation: 81 (n=1) | | | suggested automixis with central f | | 8% of males produce | Funk et al. (2010) |
| Baetidae | *Alainites (Takobia) muticus* | *Baetis pumilus* | PAL (west + east) | France | Isère | Le Charmeyran, à La | **Mixed** | fac. | balanced: 0.56 (n=43) | | (17.4-100, n=7) | 23-56d at 13.5°C | | | | | | | | Degrange (1955, 1960) |
| Baetidae | *Alainites (Takobia) muticus* | *Alainites muticus* | PAL (west + east) | Ukraine | Yasynuvats'kyi district | Bulavina River: 48°18 | **Mixed** | obl. | females only: 1.00 (n=622) | | | | | | | | | | | Martynov (2013); Alexander V. Martynov (personal communication) |
| Baetidae | *Alainites (Takobia) muticus* | | PAL (west + east) | Switzerland | | several populations | **Mixed** | mixed | from balanced to females only | | (1-98.5), 17 populati | 40.4d (25-112d at 10°C) | 55 days (21-82) | | | | | | whole Switzerland | Maud Liégeois (personal observations) |
| Baetidae | *Americabaetis sp.* | *Acerpenna sp.* | NEO | Costa Rica | Rìo Tempisquito | 10°57'25''N, 85°29'4 | **Facultative** | fac. | both sexes | | (25.8-68.5, n=7) | median=23 ±1d (n=7) | | | | | | | | Jackson&Sweeney (1995) |
| Baetidae | *Anafroptilum minor* | *Centroptilum minor* | NEA | Pennsylvania; South Carolina | | two populations | **Mixed (allopatry)** | obl. | females only | | | | | | | | clonal: suggested apomixis or automixis (without recom | | | Funk et al. (2006, 2010); Webb et al. (2012) |
| Baetidae | *Anafroptilum minor* | *Centroptilum minor* | NEA | New York | | four populations | **Mixed (allopatry)** | sex.? | both sexes | | | | | | | | | | | Funk et al. (2006, 2010); Webb et al. (2012) |
| Baetidae | *Anafroptilum semirufum* | *Centroptilum semirufum* | NEA | Pennsylvania | Chester Co. | White Clay Creek (W | **Facultative** | fac. (mixed in symp | slightly female-biased: 0.71 (n=21) | | 50 (5-75, n=5) | about twice as long as fertilised at 20°C | | forced copulation: 100 (n=1) | | | suggested automixis with central f | | 2% of males produce | Funk et al. (2010) |
| Baetidae | *Baetis (Patites) alpinus* | *Baetis dorieri* | West PAL | France | Isère | Le Charmeyran, à La | **Sexual** | sex. | both sexes | | 0 | | | | | | | | | Degrange (1960) |
| Baetidae | *Baetis (Patites) alpinus* | *Baetis alpinus* | West PAL | Czech Republic | Krasetín | Krasetínský brook | **Sexual** | | both sexes | | | | | | | | | 8+XX:XY | | Soldan&Putz (2000) |
| Baetidae | *Baetis (Rhodobaetis) bicaudatus* | | HOL (PAL + NEA) | Colorado | | Toland | **Mixed (allopatry)** | obl. | females only | | | | | | | | | | | Dodds (1923) |
| Baetidae | *Baetis (Rhodobaetis) bicaudatus* | | HOL (PAL + NEA) | North America | | several populations | **Mixed (allopatry)** | mixed | both sexual and parthenogenetic populations | | | | | | | | | | | Funk et al. (2006) |
| Baetidae | *Baetis (Rhodobaetis) bicaudatus* | | HOL (PAL + NEA) | Colorado | Gunnison Co. | | **Mixed (allopatry)** | sex. | both sexes | few hundred to over 2000 | | | | 97 (n=20) | 10-14d | | | | | Peckarsky et al. (2000) |
| Baetidae | *Baetis brunneicolor* | | NEA | Wisconsin | | | **Sexual** | sex. | | | 0 | | | | | | | | | Bergman&Hilsenhoff (1978) |
| Baetidae | *Baetis bundyae* | | NEA | Nunavut | Rankin Inlet | ten populations | **Mixed** | mixed | mainly female-biased (from balanced to females only) | | | | | | | | | | hatching length:2.5-4 | Giberson et al. (2007) |
| Baetidae | *Baetis (Rhodobaetis) foemina* | *Baetis foemina* | HOL (PAL + NEA) | Nunavut | Baffin Island | Lake Harbor | **"Obligate"** | obl. | females only | | | | | | | | | | | McDunnough (1936) |
| Baetidae | *Baetis (Rhodobaetis) foemina* | *Baetis foemina* | HOL (PAL + NEA) | Russia, Kazakhstan | | several populations | **"Obligate"** | obl. | females only | | | | | | | | | | | Kluge (1980, in russian) |
| Baetidae | *Baetis (Rhodobaetis) gemellus* | *Baetis gemellus* | West PAL | France | Isère | Le Charmeyran, à La | **Sexual** | sex. | both sexes | | 0 | | | | | | | | very closed to *B. rhoc* | Degrange (1960) |
| Baetidae | *Baetis hudsonicus* | | NEA | Canadian tundra; Manitoba | | several populations | **Mixed (allopatry)** | mixed | some populations with females only | | | | | | | | | | "a correlation with cc | McCafferty&Morihara (1979); Webb et al. (2012) |
| Baetidae | *Baetis (Rhodobaetis) noshaqensis* | *Baetis noshaqensis* | East PAL | Afghanistan | Pamir-Hindukush border | | **"Obligate"** | obl. | females only | | | | | | | | | | | Uéno (1966) |
| Baetidae | *Baetis (Rhodobaetis) rhodani* | *Baetis rhodani* | West PAL | France | Isère | Le Charmeyran, à La | **Sexual** | sex. | both sexes | | 0 | | | artificial fertilisation (failed) | | | | | | Degrange (1960) |
| Baetidae | *Baetis (Rhodobaetis) rhodani* | *Baetis rhodani* | West PAL | Germany | Hessen | Schlitz | **Sexual** | sex. | both sexes | | | | | 100% | ~28d at 8°C | 90% within 3 days at ≥15°C, withir | | 8+XX:XY | | Bohle (1969) |
| Baetidae | *Baetis (Rhodobaetis) rhodani* | *Baetis rhodani* | West PAL | France | Pyrénées-Atlantiques | Ruisseau Lissuraga | **Sexual** | sex. | both sexes | 2109 (1496-3396, n=6) | | | | 92.5 (78.7-99.1, n= | 36-46d at 7.5°C; 17-32d at 12.5°C; 13-19d at15°C; 9-15d at 17.5°C; 7-11d at 25°C | | | | | Benech 1972 |
| Baetidae | *Baetis (Rhodobaetis) rhodani* | *Baetis rhodani* | West PAL | England | | | **Sexual** | sex. | both sexes | | | | | 94.7 | 55-90d at 4-5.2°C; 20-45d at 7-10°C; 10-18d at 14-20°C | | | | | Elliott 1972 |
| Baetidae | *Baetis scambus* | | West PAL | France | Haute-Savoie | Le Fier, au pont de N | **Facultative** | fac. | both sexes | | 46.7 (n=1) | 27-257d (diapause, 16.3 to 4°C) | | | | | | | | Degrange (1960) |
| Baetidae | *Baetis (Rhodobaetis) tricaudatus* | *Baetis vagans* | NEA | Wisconsin | | | **Sexual** | sex. | | | 0 | | | | | | | | | Bergman&Hilsenhoff (1978) |
| Baetidae | *Baetis vernus* | | West PAL | France | Isère | Le Réaumont, à Réau | **Sexual** | sex. | both sexes | | 0 | | | | | | | | | Dergrange (1960) |
| Baetidae | *Baetis vernus* | | West PAL | Germany | Hessen | Schlitz | **Sexual** | sex. | both sexes | | | | | >94 | 150-270d at 6.8°C (diapause) | | | 8+XX:XY | | Bohle (1969) |
| Baetidae | *Baetis vernus* | | West PAL | Czech Republic | Kaplice | Malše River | **Sexual** | | both sexes | | | | | | | | | 8+XX:XY | | Soldan&Putz (2000) |
| Baetidae | *Bugilliesia notabilis* | *Centroptilum notabile* | AFR | Uganda | | Jinja | **Facultative** | mostly obl. | strongly female-biased: 0.97 | | has not been confirmed by hatching experiment | | | | | | | | | Tjonneland (1960) |
| Baetidae | *Bungona sp.* | *Chopralla sp.* | ORI | China | Hong Kong | | **Facultative** | fac. | both sexes | 413 (103-976, n= | 71 ±8.8 (n=4) | 11.6 ±1.1d (6-19, n=4, at 22±2°C) | | | | | | | | Salas&Dudgeon (1999) |
| Baetidae | *Callibaetis ferrugineus* | *Callibaetis americanus* | NEA | Ontario | | several populations | **Sexual** | sex.? | both sexes | | | | | | | | | | high densities, female | Ide (1930) |
| Baetidae | *Callibaetis fluctuans* | | NEA | Pennsylvania | | | **Sexual** | sex. | both sexes | | 0 | | | | | | | | ovoviviparous specie | Funk et al. (2010) |
| Baetidae | *Centroptilum lituratum* | *Centroptilum ? lituratum* | West PAL | France | Haute-Savoie | Le Fier, au pont de N | **Facultative** | fac. | both sexes | | 90.3 (n=1) | 222-265d (diapause, 15.5 to 6.5°C) | | | | | | | | Degrange (1960) |
| Baetidae | *Centroptilum luteolum* | | PAL (west + east) | France | Haute-Savoie | Lac d'Annecy | **Facultative** | fac. | balanced: 0.5 | 960-1513 | 48.7 (n=9, 3 generati | 21-94d (according to T°C) | ~1 month | 87.2 (n=1); survival | ~1% | 48 hours | | | up to 17% of males p | Degrange (1954, 1956, 1960) |
| Baetidae | *Centroptilum luteolum* | | PAL (west + east) | France | Isère | Etang de Réaumond | **Facultative** | fac. | both sexes | | 57.1 (n=30, 2 genera | 14-164d (according to T°C) | | | | | | | deuterotokous parth | Degrange (1960) |
| Baetidae | *Centroptilum sp.* | | ORI | China | Hong Kong | | **Facultative** | fac. | both sexes | 941 (204-1360, n | 41 ±6.5 (n=4) | 13.7 ±0.9d (6-28, n=4, at 22±2°C) | | | | | | | | Salas&Dudgeon (1999) |
| Baetidae | *Cloeon dipterum* | | HOL (PAL + NEA) | Sweden | | | **Sexual** | sex. | | ~700-800 | 0 | | | | | | | | ovoviviparous specie | Bengtsson (1913) |
| Baetidae | *Cloeon dipterum* | | HOL (PAL + NEA) | France | Isère | Grenoble (2 populati | **Sexual** | sex. | both sexes | 898-910 | 0 | | | hatching | | | | | ovoviviparous specie | Degrange (1960) |
| Baetidae | *Cloeon dipterum* | *Cloeon cognatum* | HOL (PAL + NEA) | Pennsylvania | | | **Sexual** | sex. | | | 0 | | | | | | | | ovoviviparous specie | Funk et al. (2010) |
| Baetidae | *Cloeon dipterum* | | HOL (PAL + NEA) | Germany, The Netherlands | | | **Sexual** | | both sexes | | | | | | | | | 8+XX:XY | ovoviviparous specie | Wolf (1960); Kiauta&Mol (1977) |
| Baetidae | *Cloeon (Similicloeon) praetextum* | *Cloeon praetextum* | West PAL | Finland | | 62°50'N, 24°50'E | **Mixed (allopatry)** | obl. | females only | | | | | | | | | | | Hirvenoja (1964) |
| Baetidae | *Cloeon (Similicloeon) praetextum* | | West PAL | Europe | | several populations | **Mixed (allopatry)** | sex.? | both sexes | | | | | | | | | | males discribed | see Ephemeroptera galactica (http://www.insecta.bio.spbu.ru/z/Eph-spp/)praetextum(Cloeon).htm) |
| Baetidae | *Cloeon (Similicloeon) simile* | *Cloeon simile* | PAL (west + east) | France | Haute-Savoie | Lac d'Annecy | **Facultative** | fac. | both sexes | 2378-3415 | ≥ 50 | 35-70d at 12°C | ~5 weeks | 1280 larvules (n=1 | 22-24d at 12°C | 48 hours | | | | Degrange (1955, 1960) |
| Baetidae | *Cloeon (Similicloeon) simile* | *Cloeon simile* | PAL (west + east) | France | Savoie | Lac de Saint André | **Facultative** | mixed (sympatry) | both sexes | | 10.9 (n=1) | 15-68d | | | | | | | thelytokous parthenc | Degrange (1960) |
| Baetidae | *Cloeon (Similicloeon) simile* | *Cloeon simile* | PAL (west + east) | France | Isère | Etang de Réaumond | **Facultative** | mixed (sympatry) | both sexes | | 8.0 (1.2-18.1, n=6) | 19-194d (according to T°C) | ~5 weeks | | | | | | thelytokous parthenc | Degrange (1960) |
| Baetidae | *Cloeon (Similicloeon) simile* | *Cloeon simile* | PAL (west + east) | Scotland | Ardamurchan Peninsul | 56°43'N, 6°13'W | **Facultative** | fac. | balanced: 0.5 (n=2261, 13 years) | | ~30 (17-35, n=7 generations) | | | | | | | | thelytokous parthenc | Harker (1997) |
| Baetidae | *Cloeon sp.* | | PAL (west + east) | France | Isère | Lac de l'Infernet, Ma | **Facultative** | fac. | both sexes | 513-991 | 26.8 (12.4-47.2, n=6) | 160-330d (diapause) | 4.5 months | artificial fertilisatic | 130d (71-171) | 2.5 months | | | thelytokous parthenc | Degrange (1960) |
| Baetidae | *Diphetor (Nigrobaetis) hageni* | *Baetis hageni* | NEA | Wisconsin | Waukesha Co. | Jericho Creek | **Mixed (allopatry)** | obl. | females only: 1.00 (n=26) | | 97.3 (97-98, n=3) | 8.5d (8-9 at 20-22°C, n=2); 19d at 13-15°C (n=1) | | | | | | | | Bergman&Hilsenhoff (1978) |
| Baetidae | *Diphetor (Nigrobaetis) hageni* | *Baetis parvus* | NEA | Ontario | | Homing's Mills | **Mixed (allopatry)** | obl. | females only | | | | | | | | | | | Ide (1930) |
| Baetidae | *Diphetor (Nigrobaetis) hageni* | *Diphetor hageni* | NEA | Pennsylvania | Chester Co. | WCC: 39°51'32''N, 7 | **Mixed (allopatry)** | obl. | females only | | | | | | | | clonal | | | Funk et al. (2006, 2008) |
| Baetidae | *Diphetor (Nigrobaetis) hageni* | *Baetis hageni* | NEA | Ontario | | Salem Creek: 43°35'N | **Mixed (allopatry)** | mostly obl. | almost females only: 0.99 (n=758) | | | | | | | | | | | Harper&Harper (1984) |
| Baetidae | *Diphetor (Nigrobaetis) hageni* | *Baetis parvus; Diphetor hage* | NEA | Colorado; Indiana, Florida | | several populations | **Mixed (allopatry)** | sex.? | both sexes | | | | | | | | | | | Dodds (1923); Funk et al. (2006); Webb et al. (2012) |
| Baetidae | *Guloptiloides gargantua* | | AFR | Madagascar | | several populations | **"Obligate"** | obl. | females only | | | | | | | | | | | Gattolliat&Sartori (2000) |
| Baetidae | *Labiobaetis frondalis* | *Baetis frondalis* | NEA | Wisconsin | Dane Co. | Dunlap Creek | **Facultative** | fac. | balanced: 0.5 | | 30.8 (10-60, n=5) | 15.5d (14-18 at 19-23°C, n=4); 31d (n=1, at 13-15°C) | | | | | | | | Bergman&Hilsenhoff (1978) |
| Baetidae | *Labiobaetis frondalis* | *Pseudocloeon frondale* | NEA | Pennsylvania | | | **Facultative** | fac. | balanced: 0.44 (n=54) | | 66 (31-92, n=14) | about twice as long as fertilised at 20°C | | forced copulation: 94 (78-99, n=6) | | | suggested automixis with central f | | thelytokous parthenc | Funk et al. (2010) |
| Baetidae | *Labiobaetis geminatus* | *Baetis geminatus* | ORI | India | | Vagai River, at Thiru | **Facultative** | fac. | balanced: 0.45 (n=94) | 930.5 (633-1237 | 32.8 (26.1-37.6, n=3) | 8-26d at 24-32°C | | | | | | | | Sivaramakrishnan et al. (1991) |
| Baetidae | *Labiobaetis propinquus* | *Baetis spinosus* | NEA | Wisconsin | Marquette Co. | Mecan River | **Facultative** | fac. | female-biased: 0.67 | | 40.4 (12-63, n=5) | 19.3d (19-20 at 19-23°C, n=4); 32d at 13-15°C (n=1) | | | | | | | | Bergman&Hilsenhoff (1978) |
| Baetidae | *Neocloeon alamance* | *Centroptilum alamance* | NEA | North Carolina | | Big Alamance Creek | **Facultative** | fac. | balanced: 0.5 (n=44) | | 75 (37-95, n=10); sur | about twice as long as fertilised | | forced copulation: 67 (25-90, n=16); survival=49% (20-72, n=15) | | | suggested automixis with central f | | not clonal; "high leve | Funk et al. (2006, 2010) |
| Baetidae | *Neocloeon triangulifer* | *Cloeon triangulifer* | NEA | Quebec | | Pond, at Rigaud | **"Obligate"** | obl. | females only: 1.00 (n >4000, 3 years) | | 86 (several females) | 12.1d (11-14 at room T°C) | 4 days | | | | | | | Gibbs (1973, 1977) |
| Baetidae | *Neocloeon triangulifer* | *Cloeon triangulifer; Centropti* | NEA | Pennsylvania | Chester Co. | White Clay Creek (W | **"Obligate"** | obl. | females only: 1.00 (severa | 1000-2000 | 87.7 (50-96, several | 7-37d at 15-25°C | 2-3 days for 80-90% | forced copulation: 33.3 (n=2) | | | clonal; suggested automixis (witho | | thelytokous parthenc | Sweeney et al. (1993); Sweeney&Vannote (1984, 1987); Funk et al. (2006) |
| Baetidae | *Neocloeon triangulifer* | *Cloeon triangulifer; Centropti* | NEA | Pennsylvania, North Carolina, South Carolina | | pop(ALA): 35.9450, - | **"Obligate"** | obl. | females only | | | | | | | | clonal | | | Sweeney et al. (1993); Funk et al. (2006); Webb et al. (2012) |
| Baetidae | *Nigrobaetis niger* | *Baetis niger* | West PAL | France | Isère | Le Méaudret, à Méa | **Mixed (allopatry)** | fac. | both sexes | | 35.5 (n=1) | 18-40d at 15.5°C | | | | | | | | Degrange (1960) |
| Baetidae | *Nigrobaetis niger* | | West PAL | Crimea | | Mountain region | **Mixed (allopatry)** | obl. | females only | | | | | | | | | | | Martynov (2013) |
| Baetidae | *Procloeon fragile* | | NEA | Pennsylvania | Chester Co. | WCC: 39.86061°N, 7 | **Facultative** | fac. | balanced: 0.41 (n=12) | | 40 (30-57, n=6); surv | about twice as long as fertilised at 20°C | | forced copulation: 84 (70-99, n=6); survival: 39% (14-79, n=6) | | | suggested automixis with central f | | thelytokous parthenc | Funk et al. (2010) |
| Baetidae | *Procloeon pennulatum* | *Centroptilum pennulatum* | HOL (PAL + NEA) | France | Isère | Le Méaudret, à Méa | **Facultative** | fac. | both sexes | 1327-2410 | (20.3-44.5, n=18); dv | 18-307d at 16.5 to 6.5°C (partial diapause) | | | | | | | | Degrange (1960) |
| Baetidae | *Procloeon rivulare* | | NEA | Pennsylvania | Chester Co. | WCC: 39.86061°N, 7 | **Facultative** | fac. | balanced: 0.4 (n=67) | | 45 (11-82, n=14); sur | about twice as long as fertilised at 20°C | | forced copulation: 69 (16-97, n=9); survival: 16% (0-34, n=7) | | | suggested automixis with central f | | thelytokous parthenc | Funk et al. (2010) |
| Baetidae | *Procloeon rufostrigatum* | *Centroptilum rufostrigatum* | NEA | Eastern North America | | | **Sexual** | tycho. | both sexes | | <0.1 (0-<0.1) | | | | | | | | | Sweeney&Vannote (1987) |
| Baetidae | *Procloeon viridoculare* | *Procloeon irrubrum* | NEA | Manitoba | | several populations | **Mixed (allopatry)** | obl. | females only | | | | | | | | | | | Lowen&Flannagan (1992) |
| Baetidae | *Procloeon viridoculare* | | NEA | South Carolina | Edgefield Co. | Turkey Creek: 33.79 | **Mixed (allopatry)** | obl. | females only | | | | | | | | clonal: suggested apomixis or autc | | "the genetic variatior | Funk et al. (2008, 2010); Webb et al. (2012) |
| Baetidae | *Procloeon viridoculare* | *Centroptilum viridocularis; Pr* | NEA | Florida; Eastern US | | | **Mixed (allopatry)** | sex.? | both sexes | | | | | | | | | | | Berner (1940); Funk et al. (2010) |
| Baetidae | *Procloeon sp.* | | ORI | China | Hong Kong | | **Facultative** | fac. | both sexes | 1547 (n=1) | 73 (n=1) | 13.2d (9-21 at 22 ±2°C) | | | | | | | | Salas&Dudgeon (1999) |
| Baetiscidae | *Baetisca rogersi* | | NEA | Florida | Gadsden Co. | | **Sexual** | tycho. | balanced: 0.52 (n=402, 2 | 2168 (1500-2800 | 1 (0.3-1.6, n=5) | 1st hatch: 26.6d (20-31, n=5) | | forced copulation: | 1st hatch: 21d (20-23, n=5) | | | | | Pescador (1973); Perscador&Peters (1974) |
| Behningiidae | *Dolania americana* | | NEA | Florida | Okaloosa Co., Santa Ro | Blackwater River | **Sexual** | sex. | balanced | 91 (24-161, n=82 | 0 | | | | | | | | "females never molt i | Peters&Peters (1977) |
| Behningiidae | *Dolania americana* | | NEA | South Carolina | | Upper Three Runs: 3 | **Sexual** | sex. | balanced: 0.5 | 77 (44-148, n=18) | | | | | diapause: ~10 months | | | | "females never molt i | Harvey et al. (1980); Sweeney&Vannote (1982) |
| Caenidae | *Caenis catherinae* | | NEO | Gadeloupe | Basse Terre | several populations | **"Obligate"** | obl. | females only | | | | | | | | | | | Hofmann et al. (1999) |
| Caenidae | *Caenis cuniana* | | NEO | Brazil | State of São Paulo | Boracéia | **"Obligate"** | obl. | females only: 1.00 | ~250 (100-400) | 87.4 (72.3-99.4, n=7) | 18.2d (14-23 at 21°C, n=1); 11.3d (9-25 at 23.5°C, n=2); 10.4d (7-29 at 25.5°C, n=2) | | | | | | | thelytokous parthenc | Froehlich (1969) |
| Caenidae | *Caenis cuniana* | | NEO | Brazil | State of Rio de Janeiro | Restinga de Barra de | **"Obligate"** | obl. | females only: 1.00 | | 71 (62-78, n=3) | 11.6d (9-25 at 22°C, n=3) | | | | | | | thelytokous parthenc | Da-Silva (1993, 1998) |
| Caenidae | *Caenis femina* | | NEO | Guadeloupe | Basse Terre | several populations | **"Obligate"** | obl. | females only | | | | | | | | | | | Hofmann et al. (1999); Michel Sartori (personal communication) |
| Caenidae | *Caenis horaria* | | PAL (west + east) | France | Isère | Etang, à Réaumont | **Sexual** | sex. | both sexes | 505-607 | 0 | | | | | | | | | Degrange (1960) |
| Caenidae | *Caenis horaria* | *Caenis dimidiata* | PAL (west + east) | Europe | | | **Sexual** | tycho.? | | | hatching | | | | | | | | "un cas de parthénog | G. Zerega-Fombona (unpublished data) |
| Caenidae | *Caenis horaria* | | PAL (west + east) | The Netherlands | Utrecht | | **Sexual** | | both sexes | | | | | | | | | 6+XX:XO | | Mol (1978) |
| Caenidae | *Caenis knowlesi* | | AFR | Gabon | Moyen-Ogooué | Kéri (Lambaréné) | **"Obligate"** | obl. | females only: 1.00 | 526 ±151 (271-7 | 93 (88.2-99.4, n=5) | 1st hatch: 7-10d at 25-26°C | | | | | | | only one male discrib | Gillies&Knowles (1990) |
| Caenidae | *Caenis luctuosa* | *Caenis moesta* | PAL (west + east) | France | Isère | Lac de Saint-Sixte | **Sexual** | tycho. (mixed) | both sexes | 765-1103 | (1-10.8) | 15-36d at 16°C | | | | | | | | Degrange (1960) |
| Caenidae | *Caenis macrura* | | PAL (west + east) | France | Isère, Drôme | | **Sexual** | sex. | both sexes | 675 | 0 | | | | | | | | | Degrange (1960) |
| Caenidae | *Caenis macrura* | | PAL (west + east) | Czech Republic | Holubov | Křemžský brook | **Sexual** | | both sexes | | | | | | | | | 6+XX:XO | | Soldan&Putz (2000) |
| Caenidae | *Caenis robusta* | | PAL (west + east) | France | Isère | Grenoble | **Sexual** | sex. | both sexes | | 0 | | | | | | | | | Degrange (1960) |
| Caenidae | *Caenis vanuatensis* | | PAC | Vanuatu | Espirito Santo Island | several populations | **"Obligate"** | obl. | females only | | | | | | | | | | | Malzacher&Staniczek (2007) |
| Dipteromimidae | *Dipteromimus tipuliformis* | | East PAL | Japan | | | **Sexual** | sex. | both sexes | | 0 (n=6) | | | forced copulation: 91.5 ±6.0 (n=18); natural mating: 91.5 ±13.2 (n=7) | | | | | | Takenaka et al. 2019 |
| Ephemerellidae | *Ephemerella aurivillii* | *Chitonophora aurivillii* | HOL (PAL + NEA) | Lithuania, Latvia, Estonia | | several populations | **Mixed (allopatry)** | obl. | females only | | | | | | | | | | | Kazlauskas (1963) |
| Ephemerellidae | *Ephemerella aurivillii* | *Ephemerella taeniata* | HOL (PAL + NEA) | Scandinavia, Alaï | | several populations | **Mixed (allopatry)** | sex.? | both sexes | | | | | | | | | | | Tshernova (1952) *in* Kazlauskas (1963) |
| Ephemerellidae | *Ephemerella aurivillii* | | HOL (PAL + NEA) | New York | Essex Co. | 44.416, -73.911 | **Mixed (allopatry)** | sex.? | both sexes | | | | | | | | | | | Webb et al. (2012) |
| Ephemerellidae | *Ephemerella dorothea* | | NEA | Eastern North America | | | **Sexual** | tycho. | both sexes | | <0.1 (0-<0.1) | | | | | | | | | Sweeney&Vannote (1987) |
| Ephemerellidae | *Ephemerella invaria* | *Ephemerella feminina* | NEA | New York | | near McLean | **Mixed (allopatry)** | obl. | females only | | | | | | | | | | | Needham (1924) |
| Ephemerellidae | *Ephemerella invaria* | | NEA | Quebec | | several populations | **Mixed (allopatry)** | mostly obl. | strongly female-biased: 0.93 (n=180) | | | | | | | | | | "There is some indica | Harper&Harper (1982) |

| | | | | | | | | | | | | | | | | | | | | |
|---|---|---|---|---|---|---|---|---|---|---|---|---|---|---|---|---|---|---|---|---|
| Ephemerellidae | *Ephemerella invaria* | | NEA | New York, Pennsylvania, Tennessee | | several populations | **Mixed (allopatry)** | tycho.? | both sexes | | hatching | | | | | | | | "bisexual mayfly popu | Funk et al. (2010); Webb et al. (2012) |
| Ephemerellidae | *Ephemerella sp.* | | NEA | Colorado | | | | obl. | females only | | | | | | | | | | | Dodds (1923) |
| Ephemerellidae | *Ephemerella mucronata* | *Ephemerella (Chitonophora)* | HOL (PAL + NEA) | France | Haute-Savoie | Le Fier, au pont de N | **Facultative** | fac. (mixed in symp | both sexes | 1230-1671 | (4.4-72.5, n=10); mo | 32d (26-76 at 14-18°C) | | | | | | | | Degrange (1960) |
| Ephemerellidae | *Ephemerella mucronata* | | HOL (PAL + NEA) | Czech Republic | Krasetín | Krasetínský brook | **Facultative** | | both sexes | | | | | | | | | 6+XX:XY | | Soldan&Putz (2000) |
| Ephemerellidae | *Ephemerella notata* | | West PAL | Czech Republic; central | Poland | several populations | **Mixed** | obl. | females only | | | | | | | | | | geographical parthen | reviewed in Glazaczow (2001) |
| Ephemerellidae | *Ephemerella notata* | | West PAL | northern Poland | | several populations | **Mixed** | mixed | from balanced to female-biased | | | | | | | | | | geographical parthen | reviewed in Glazaczow (2001) |
| Ephemerellidae | *Ephemerella notata* | | West PAL | England, Danmark | | several populations | **Mixed** | sex.? | both sexes | | | | | | | | | | geographical parthen | reviewed in Glazaczow (2001) |
| Ephemerellidae | *Eurylophella doris* | | NEA | Eastern US | | several populations | **Sexual** | tycho.? | both sexes | | hatching | | | | | | | | "bisexual mayfly popu | Funk et al. (2010); Webb et al. (2012) |
| Ephemerellidae | *Eurylophella enoensis* | | NEA | North Carolina | | Coon Creek: 36.354, | **Sexual** | tycho.? | both sexes | | hatching | | | | | | | | "bisexual mayfly popu | Funk et al. (2010); Webb et al. (2012) |
| Ephemerellidae | *Eurylophella funeralis* | *Ephemerella funeralis* | NEA | New Hampshire | White Mountains | Hubbard brook wate | **Mixed** | fac. | female-biased: 0.89 (n=19 | 1850 ±87 | 53 (n=1) | 32d at room T°C | | | | | | | several populations | Fiance (1978) |
| Ephemerellidae | *Eurylophella funeralis* | | NEA | Quebec | | tributary of the Mata | **Mixed** | obl. | females only: 1.00 (n=9) | | 86.7 ±11.7 (65.9-97.( | 36.1 ±2.2d (34-39, n=8, at 14.5±1°C) | | | | | | | geographical parthen | Sweeney&Vannote (1987) |
| Ephemerellidae | *Eurylophella funeralis* | | NEA | Quebec | | tributary of the Mata | **Mixed** | obl. | females only: 1.00 (n=37) | | 83.4 ±24.3 (17.7-97.: | 34.7 ±1.8d (31-38, n=30 at 14.5±1°C) | | | | | | | geographical parthen | Sweeney&Vannote (1987) |
| Ephemerellidae | *Eurylophella funeralis* | | NEA | Delaware | New Castle Co. | White Clay Creek (DE | **Mixed** | obl. | females only: 1.00 (n=52) | | 80.9 ±14.5 (54.2-98.4 | 33.7 ±1.9d (30-39, n=50 at 14.5±1°C) | | | | | | | geographical parthen | Sweeney&Vannote (1987) |
| Ephemerellidae | *Eurylophella funeralis* | | NEA | South Carolina | Oconee Co. | Cranes Creek (SC) | **Mixed** | obl. | females only: 1.00 (n=9) | | 88.5 ±10 (74-97.1, n= | 37.4 ±2.4d (33-41, n=9 at 14.5°C) | | | | | | | geographical parthen | Sweeney&Vannote (1987) |
| Ephemerellidae | *Eurylophella funeralis* | | NEA | Pennsylvania | Chester Co. | White Clay Creek (PA | **Mixed** | obl. | females only: 1.00 (n=27) | | 84.9 ±15.1 (48-97.5, | 34.0 ±2.4d (29-38, n=27 at 14.5±1°C) | | artificial fertilisation | | | clonal: suggested apomixis or auto | | thelytokous parthenc | Sweeney&Vannote (1987) |
| Ephemerellidae | *Eurylophella funeralis* | | NEA | Pennsylvania | Chester Co. | White Clay Creek (PA | **Mixed** | obl. | females only | | | 27.7 ±1d (27-29, n=5 at 12-19°C) | | | | | | | | Sweeney&Vannote (1987) |
| Ephemerellidae | *Eurylophella funeralis* | | NEA | Pennsylvania | Chester Co. | White Clay Creek (PA | **Mixed** | obl. | females only | | | 26.5 ±0.6d (26-28, n=15 at 12-19°C) | | | | | | | | Sweeney&Vannote (1987) |
| Ephemerellidae | *Eurylophella funeralis* | | NEA | Pennsylvania | Chester Co. | tributary to Brandyw | **Mixed** | obl. | females only | | | 26.7 ±0.6d (26-28, n=11 at 12-19°C) | | | | | | | | Sweeney&Vannote (1987) |
| Ephemerellidae | *Eurylophella funeralis* | | NEA | New Hampshire | Grafton Co. | tributary of Hubbard | **Mixed** | mostly obl. | almost females only: 0.98 (n=38) | | 86.8 ±9.5 (66.9-97.9, | 38.1 ±4.2d (30-53, n=26 at 14.5 ±1°C) | | | | | | | | Sweeney&Vannote (1987) |
| Ephemerellidae | *Eurylophella funeralis* | | NEA | Vermont | Bennington Co. | Tributary of the Batt | **Mixed** | fac. | female-biased: 0.87 (n=17) | | 62.8 ±25.2 (23.2-87.4 | 36.9 ±3.6d (32-40, n=6 at 14.5°C) | | | | | | | | Sweeney&Vannote (1987) |
| Ephemerellidae | *Eurylophella funeralis* | | NEA | Ohio | Lawrence Co. | Storms Creek (OH) | **Mixed** | fac. (mixed in symp | balanced: 0.5 | | 18.5 ±15.9 (n=9) | 34.3 ±3.7d (29-37, n=4 at 12-19°C) | | artificial fertilisation | | | | | | Sweeney&Vannote (1987) |
| Ephemerellidae | *Eurylophella funeralis* | | NEA | West Virginia | Cabell Co. | Hisey Fork of Four Po | **Mixed** | fac. (mixed in symp | balanced: 0.5 | 2227 (1709-3069 | 14.2 ±15.1 (n=25) | 32.5 ±3.1d (28-40, n=15 at 12-19°C) | | artificial fertilisation | | | | | | Hamilton&Tarter (1977); Sweeney&Vannote (1987) |
| Ephemerellidae | *Eurylophella funeralis* | | NEA | Pennsylvania | Susquehanna Co. | Nine Partners Creek | **Mixed** | tycho. (mixed) | balanced: 0.5 (n=35) | | 6.1 ±5.2 (0.3-15.8, n= | 44.2 ±4.5d (36-57, n=39 at 14.5°C) | | | | | | | | Sweeney&Vannote (1987) |
| Ephemerellidae | *Eurylophella funeralis* | | NEA | Ontario, Georgia, Pennsylvania, Virginia, Quebe | | several populations | **Mixed** | obl. | females only | | | | | | | | | | | Sweeney&Vannote (1987); Webb et al. (2012) |
| Ephemerellidae | *Eurylophella funeralis* | | NEA | Virginia, New York, Pennsylvania, Connecticut, | | several populations | **Mixed** | sex.? | both sexes | | | | | | | | | | | Sweeney&Vannote (1987); Webb et al. (2012) |
| Ephemerellidae | *Eurylophella oviruptis* | | NEA | North Carolina | Robeson Co. | Back Swamp: 34.600 | **"Obligate"** | obl. | females only: 1.00 | 1450 (n=1) | 79 (21-98, n=15) | 1st hatch: 43d (40-47, n=7 at 5.6-17.8°C); 23d (17-29, n=8 at 16°C) | | | | | probably clonal | | Heterozygosity: 0.08 | Funk et al. (2008) |
| Ephemerellidae | *Eurylophella oviruptis* | | NEA | North Carolina | | several populations | **"Obligate"** | obl. | females only | | | | | | | | | | | Funk et al. (2008) |
| Ephemerellidae | *Serratella (Torleya) ignita* | *Ephemerella ignita* | PAL (west + east) | France | | three populations | **Mixed (sympatry)** | mixed (sympatry) | both sexes | 842-1171 | (1.4-25.6) | 185-279d at 16.6-7°C | | forced copulation (failed) | | | | | | Degrange (1960) |
| Ephemerellidae | *Serratella (Torleya) ignita* | *Ephemerella ignita* | PAL (west + east) | England | Lake District | Wilfin Beck | **Mixed (sympatry)** | sex.? | both sexes | 148-603 | | | | >85 | 112-392d at 8°C; 56-266d at 10°C; 56-224d at 12°C; 56-196d at 14°C; 56-294d at 16°C | | | | | Elliott (1978) |
| Ephemerellidae | *Serratella (Torleya) ignita* | *Ephemerella ignita* | PAL (west + east) | Belgium | Malmédy | | **Mixed (sympatry)** | | both sexes | | | | | | | | | 14+XX:XY | | Mol (1978) |
| Ephemerellidae | *Serratella sp.* | | NEA | Colorado | | Frying Pan River | | | | | | | | | | | | | "presumably parthen | G. F. Edmunds and M. Lawson (unpublished) *in* Funk et al. (2008) |
| Ephemerellidae | *Torleya major* | *Ephemerella (Torleya) belgica* | West PAL | France | Isère | Ruisseau de St-Etien | **Sexual** | tycho. | both sexes | 1689 | 0.8 (n=1) | 39-50d at 13.4-16.6°C | | | | | | | | Degrange (1960) |
| Ephemerellidae | *Torleya major* | *Ephemerella (Torleya) belgica* | West PAL | France | Drôme | L'Herbasse, à Pont-d | **Sexual** | tycho. | both sexes | 730 | 1.4 (n=1) | 34-46d at 15.7-16.6°C | | | | | | | | Degrange (1960) |
| Ephemeridae | *Ephemera danica* | | West PAL | France | Ardèche, Isère, Savoie | three populations | **Sexual** | sex. | both sexes | 2434-4600 | 0 | | | artificial fertilisation: 0.4-1.5 | | | | | sometimes, females r | Degrange (1960) |
| Ephemeridae | *Ephemera danica* | | West PAL | England | Hampshire | | **Sexual** | sex. | both sexes | 3000-6000 | | | | 90 | | | | | | Bennett (2007) |
| Ephemeridae | *Ephemera danica* | | West PAL | Germany, Luxembourg, Czech Republic | | several populations | **Sexual** | | both sexes | | | | | | | | | 10+XX:XO | | Wolf (1946); Mol (1978); Soldan&Putz (2000) |
| Ephemeridae | *Ephemera (Sinephemera) glaucops* | *Ephemera glaucops* | West PAL | France | Haute-Savoie | Lac d'Annecy | **Sexual** | sex. | both sexes | 2787 | 0 | | | | | | | | | Degrange (1960) |
| Ephemeridae | *Ephemera (Sinephemera) japonica* | *Ephemera japonica* | East PAL | Japan | Nagano Prefecture: Ma | two populations | **Sexual** | tycho. | both sexes | 1900.83 (n=12) | 1.1 ±1.4 (0-4.9, n=11) | | | 78.1 (n=1) | | | | 10+XX:XO | | Sekine&Tojo (2010)b |
| Ephemeridae | *Ephemera orientalis* | | East PAL | Japan | Hyōgo Prefercture: Ibo | Ibo-gawa River: 34°5 | **Sexual** | tycho. | both sexes | 3814 (n=7) | 0.4 ±0.4 (0-1.2, n=7) | | | | | | | 10+XX:XO | | Sekine&Tojo (2010)b |
| Ephemeridae | *Ephemera simulans* | | NEA | Eastern North America | Lake Erie | several populations | **Sexual** | sex. | balanced: 0.54 | 4500 (3202-5879 | 0 | | | | ~20-30d (at 18-20°C) | | | | | Britt (1962) |
| Ephemeridae | *Ephemera simulans* | | NEA | Ohio | | | **Sexual** | sex.? | both sexes | | | | | artificial fertilisatic | 1st hatch: 8d | | | | | Wiebe (1925) |
| Ephemeridae | *Ephemera (Sinephemera) strigata* | *Ephemera strigata* | East PAL | Japan | Fukushima Prefecture: | Kajiya-gawa River: 3 | **Sexual** | tycho. | both sexes | 2322.6 (n=7) | 0.7 ±1.6 (0-4.4, n=7) | | | | | | | 10+XX:XO | | Sekine&Tojo (2010)b |
| Ephemeridae | *Ephemera (Sinephemera) strigata* | *Ephemera strigata* | East PAL | Japan | Nara Prefecture | several populations | **Sexual** | sex. | both sexes | | | | | >95 (n=4) | <10d at 15-20°C;20-25d at 20°C; 77-82d at 4°C | | | | | Ban&Kawai (1986) |
| Ephemeridae | *Ephemera varia* | | NEA | Eastern North America | | | **Mixed** | obl. | females only: 1.00 | | 99 | | | | | | | | | Sweeney&Vannote (1987) |
| Ephemeridae | *Ephemera varia* | | NEA | Eastern North America | | | **Mixed** | fac. (mixed in symp | both sexes | | 10 (0.7-42) | | | | | | | | | Sweeney&Vannote (1987) |
| Ephemeridae | *Ephemera varia* | | NEA | Pennsylvania | Chester Co. | White Clay Creek (W | **Mixed** | obl. | females only | | | | | | | | clonal: suggested apomixis or automixis (without recom | | | Funk et al. (2006, 2008, 2010) |
| Ephemeridae | *Ephemera vulgata* | | PAL (west + east) | France | Haute-Savoie, Isère | four populations | **Sexual** | sex. | both sexes | | 0 | | | | | | | | sometimes, females r | Degrange (1960) |
| Ephemeridae | *Ephemera vulgata* | | PAL (west + east) | Czech Republic | Kokořín | Pšovka brook | **Sexual** | | both sexes | | | | | | | | | 12+XX:XO | | Soldan&Putz (2000) |
| Ephemeridae | *Hexagenia bilineata* | | NEA | Minnesota | Winona | Mississipi River | **Sexual** | sex. | both sexes | 2500 (380-6300, | 0 (n=12) | | | "artificially inseminated hatched readily" | | | | | | Thomforde&Fremling (1968) |
| Ephemeridae | *Hexagenia limbata* | | NEA | Michigan | | Pine Lake | **Sexual** | tycho. | balanced: 0.53 | 3005.6 (2260-35 | 0 (but some embryonic dvpmt) | | | 98.1 (95.7-99.5, n= | 11-14d at 24-35°C; 10-21d at 25°C | | | | | Hunt (1951, 1953) |
| Ephemeridae | *Hexagenia limbata* | | NEA | Michigan | | Gun Lake | **Sexual** | tycho. | balanced: 0.51 | 5430.3 (3507-76 | 0 (but some embryonic dvpmt) | | | 93.3 (91.2-96.6, n= | 20-26d at 16.5-23°C; 17-19d at 18-23°C | | | | | Hunt (1951, 1953) |
| Ephemeridae | *Hexagenia limbata* | | NEA | Michigan | | Big Silver Lake | **Sexual** | tycho. | balanced: 0.49 | 4484.3 (3831-56 | 0 (but some embryonic dvpmt) | | | 97.3 (96.3-98.5, n= | 18-22d at 19-27°C; 70d at 2-4.5°C | | | | | Hunt (1951, 1953) |
| Ephemeridae | *Hexagenia limbata* | *Hexagenia limbata occulta* | NEA | Manitoba | | Lake Winnipeg | **Sexual** | sex. | female-biased: 0.54 | 3509.5 (3388-36 | 0 (n=1) | | | artificial fertilisatic | 17-19d at 18-23°C | | | | | Neave (1932) |
| Ephemeridae | *Hexagenia limbata* | | NEA | Eastern US | | | **Sexual** | tycho.? | both sexes | | hatching | | | | | | | | "bisexual mayfly popu | Funk et al. (2010) |
| Ephemeridae | *Hexagenia rigida* | | NEA | Manitoba | Winnipeg | Red River | **Sexual** | tycho. | both sexes | 1577 (n=1) | ~4 (0-9, n=35) | | | 93.8 | 6d at 24-32°C; 23d at 20°( | 90% within 4 days at 20-32°C | | | | Friesen&Flannagan (1976); Friesen et al. (1979) |
| Heptageniidae | *Afronurus sp.* | *Cinygmina sp.* | ORI | China | Hong Kong | | **Sexual** | sex. | | | 0 | | | | | | | | | Salas&Dudgeon (1999) |
| Heptageniidae | *Dactogenia coerulans* | *Heptagenia coerulans* | West PAL | France | Drôme | Le Rhône, à Valence | **Sexual** | sex. | both sexes | | 0 | | | | | | | | | Degrange (1960) |
| Heptageniidae | *Ecdyonurus dispar* | *Ecdyonurus fluminum* | West PAL | France | Ardèche | L'Eyrieux, à Beaucha | **Sexual** | tycho. | both sexes | 1603-1994 | 0.3 (n=1) | 45-56d at 13°C | | | | | | | | Degrange (1955, 1960) |
| Heptageniidae | *Ecdyonurus dispar* | | West PAL | England | | Lake Ennerdale | **Sexual** | sex. | both sexes | | 0 (n=2) | | | artificial fertilisatic | 13-35d at 14.8-20°C | | | | | Humpesh (1980)a & b |
| Heptageniidae | *Ecdyonurus dispar* | | West PAL | England | | Lake Windermere | **Sexual** | tycho. | both sexes | | <1 (n=7) | 21-33d at 20.2°C | | artificial fertilisatic | 12-45d at 13.7-20.5°C | | | | | Humpesh (1980)a & b |
| Heptageniidae | *Ecdyonurus dispar* | | West PAL | England | | River Lune | **Sexual** | tycho. | both sexes | 2066 (n=1) | <1 (n=4) | 35-64d at 20.1°C; 74-276d at 14.1°C | | artificial fertilisatic | 18-364d at 14-20.5°C | | | | | Humpesh (1980)a & b |
| Heptageniidae | *Ecdyonurus dispar* | | West PAL | Luxembourg | Diekirch | | **Sexual** | | both sexes | | | | | | | | | 18+XX:XY | | Mol (1978) |
| Heptageniidae | *Ecdyonurus forcipula* | | West PAL | France | Isère | Ruisseau de St-Etien | **Sexual** | tycho. | both sexes | 9832 (n=1) | 0.2 (n=1) | 35-40d at 15.5°C | | | | | | | | Degrange (1960) |
| Heptageniidae | *Ecdyonurus forcipula* | | West PAL | France | Haute-Savoie | Le Fier, à Alex | **Sexual** | sex. | both sexes | 1222 (n=1) | | | | natural fertilisatio | 17-27d at 16.5°C | | | | mated female catche | Degrange (1960) |
| Heptageniidae | *Ecdyonurus (Helvetoraeticus) heleticus* | *Ecdyonurus helveticus* | West PAL | France | Drôme | Source de l'Archiane | **Sexual** | sex. | both sexes | 3973-4472 | 0 | | | ~100% (n=3) | 25-168d | 143 days (5 months) | | | | Degrange (1960) |
| Heptageniidae | *Ecdyonurus insignis* | | West PAL | France | Ardèche | L'Eyrieux, à Beaucha | **Sexual** | tycho. | both sexes | | 0.4 (0-5) | | | | | | | | | Degrange (1955, 1960) |
| Heptageniidae | *Ecdyonurus insignis* | | West PAL | England | | River Eden | **Sexual** | tycho. | both sexes | 2000-3500 | 2.7 (0.5-5.7, n=3) | 16-118d at 8.7-20°C | | artificial fertilisatic | 10-80d at 8.7-20°C | | | | | Humpesch (1980)a & b |
| Heptageniidae | *Ecdyonurus (Helvetoraeticus) picteti* | *Ecdyonurus picteti* | West PAL | Austria | | Herrnalmbach | **Sexual** | sex. | both sexes | | 0 (n=3) | | | artificial fertilisatic | 70-285d at 4-8°C; 35-64d at 10-13°C; 23-35d at 15-17°C | | | | | Humpesch (1980)a & b; Humpesch (1982) |
| Heptageniidae | *Ecdyonurus (Helvetoraeticus) picteti* | *Ecdyonurus picteti* | West PAL | Austria | | Seebach | **Sexual** | tycho. (mixed) | both sexes | up to 6900 | 1.2 (0-11.3, n=28) | 127-295d at 4.5-8°C; 60-106d at 10°C; 32-82d at | | artificial fertilisatic | 55-136d at 4.5-8°C; 35-63d at 10-13°C; 13-38d at 14.5-20°C | | | | | Humpesch (1980)a & b; Humpesch (1982) |
| Heptageniidae | *Ecdyonurus torrentis* | | West PAL | England | | River Lune | **Facultative** | fac. (mixed in symp | both sexes | up to 8000 | 20.9 (0.4-39, n=4) | 63-300d at 4-9°C; 17-56d at 14.5-20°C | | artificial fertilisatic | 46-350d at 4-9°C; 18-54d at 11-15°C; 12-24d at 17.5-20°C | | | | | Humpesch (1980)a & b |
| Heptageniidae | *Ecdyonurus torrentis* | | West PAL | Czech Republic | Třísov | Vltava River | **Facultative** | | both sexes | | | | | | | | | 18+XX:XY | | Soldan&Putz (2000) |
| Heptageniidae | *Ecdyonurus venosus* | | West PAL | England | | River Brathay | **Sexual** | sex. | both sexes | | 0 (n=4) | | | artificial fertilisatic | 53-310d at 4-9°C; 21-44d at 11-15°C; 11-20d at 18-20°C | | | | | Humpesch (1980)a & b |
| Heptageniidae | *Ecdyonurus venosus* | | West PAL | Austria | | Seebach | **Sexual** | tycho. (mixed) | both sexes | up to 6000 | 0.4 (0-11.3, n=60) | 147-391d at 4-8°C; 56-230d at 10-15°C; 16-133d | | artificial fertilisatic | 68-350d at 3.5-8°C; 32-70d at 10-13°C; 13-44d at 15-20°C | | | | | Humpesch (1980)a & b; Humpesch (1982) |
| Heptageniidae | *Ecdyonurus venosus* | | West PAL | Switzerland | Vaud | 46.52016, 6.67067 | **Sexual** | tycho. | balanced: 0.52 (n=21) | up to 3640 | 0.1 (n=1) | | | | | | | | | Maud Liégeois (personal observations) |
| Heptageniidae | *Ecdyonurus venosus* | | West PAL | Switzerland | Vaud | 46.66858, 6.69623 | **Sexual** | tycho. | balanced: 0.53 (n=19) | | 0.7 (n=1) | | | | | | | | | Maud Liégeois (personal observations) |
| Heptageniidae | *Ecdyonurus venosus* | | West PAL | Switzerland | Vaud | 46.42873, 6.19307 | **Sexual** | tycho. | balanced: 0.58 (n=12) | up to 4215 | 1.5 (n=1) | | | | | | | | | Maud Liégeois (personal observations) |
| Heptageniidae | *Ecdyonurus venosus* | | West PAL | Switzerland | Vaud | 46.82108, 6.62107 | **Sexual** | tycho. | balanced: 0.6 (n=10) | up to 3510 | 5.3 (n=2) | | | | | | | | | Maud Liégeois (personal observations) |
| Heptageniidae | *Ecdyonurus venosus* | | West PAL | Switzerland | Vaud | 46.499427, 6.38885 | **Sexual** | tycho. | both sexes | up to 1900 | 0.5 (n=1) | | | | | | | | | Maud Liégeois (personal observations) |
| Heptageniidae | *Electrogena lateralis* | *Heptagenia lateralis* | West PAL | France | Isère | Le Charmeyran, à La | **Sexual** | tycho. | both sexes | 1672-2356 | 4.5 (n=1) | 46-71d at 15.5°C | | | | | | | | Degrange (1955, 1960) |
| Heptageniidae | *Electrogena lateralis* | *Heptagenia lateralis* | West PAL | France | Haute-savoie | Le Fier, au pont de N | **Sexual** | | both sexes | | | | | forced copulation: | natural fertilisation: 17-23d at 15.5°C | | | | | Degrange (1960) |
| Heptageniidae | *Electrogena sp.* | | ORI | China | Hong Kong | | **Sexual** | sex. | | | 0 | | | | | | | | | Salas&Dudgeon (1999) |
| Heptageniidae | *Epeorus (Ironopsis) alpicola* | *Epeorus alpicola* | West PAL | France | Isère | Ruisseau de Theys | **Sexual** | tycho. | both sexes | | 4.1 (n=2); more for d | 33-43d at 15-19°C | | | | | | | | Degrange (1960) |
| Heptageniidae | *Epeorus assimilis* | | West PAL | France | Ardèche | L'Eyrieux, à Beaucha | **Sexual** | tycho. | both sexes | 3545-9341 | 0.9 (n=1) | 22-36d at 15°C | | | | | | | | Degrange (1955, 1960) |
| Heptageniidae | *Epeorus assimilis* | | West PAL | France | Isère | Ruisseau de St-Etien | **Sexual** | sex.? | both sexes | 210 | | | | 69.5 (n=1, but som | 17d at 14°C | | | | mated female catche | Degrange (1960) |
| Heptageniidae | *Heptagenia sulphurea* | | PAL (west + east) | France | Drôme | Le Rhône, à Valence | **Sexual** | tycho. | both sexes | | 0.5 (n=1) | 205-207d at 10°C | | | | | | | | Degrange (1960) |
| Heptageniidae | *Heptagenia sulphurea* | | PAL (west + east) | France | Haute-Savoie | Lac d'Annecy | **Sexual** | sex.? | both sexes | | | | | >80 (n=1) | 28-206d (facultative diapause) | | | | | Degrange (1960) |
| Heptageniidae | *Heptagenia sulphurea* | | PAL (west + east) | Czech Republic | Kaplice | Malše River | **Sexual** | | both sexes | | | | | | | | | 18+XX:XY | | Soldan&Putz (2000) |
| Heptageniidae | *Maccaffertium mediopunctatum* | *Stenonema nepotellum* | NEA | Indiana | | | **Sexual** | sex. | | | 0 | | | | | | | | | McCafferty&Huff (1974) |
| Heptageniidae | *Maccafertium mexicanum* | *Maccafertium integrum* | PANA (NEA + NEO) | Eastern US | | | **Sexual** | tycho.? | | | hatching | | | | | | | | "species that we had | Funck et al. (2010) |
| Heptageniidae | *Maccafertium modestum* | | NEA | Esatern US | | | **Sexual** | tycho.? | | | hatching | | | | | | | | "species that we had | Funck et al. (2010) |
| Heptageniidae | *Maccafertium modestum* | *Stenonema modestum* | NEA | Virginia | | North Anna River | **Sexual** | | female-biased: 0.66 (n=2! | 473-515 (n=4) | | | | ~20 (but ovipositer | 25-28d | | | | "fungal contaminatio | Kondratieff&Voshell (1980) |
| Heptageniidae | *Maccaffertium pulchellum* | *Stenonema pulchellum* | NEA | Indiana | | | **Sexual** | tycho. | both sexes | 3483 (n=1) | 0.3 (n=1) | 14-17d at 18°C | | | | | | | | Huff&McCafferty (1974) |
| Heptageniidae | *Maccaffertium vicarium* | *Stenonema vicarium* | NEA | Indiana | | | **Sexual** | tycho. | both sexes | 3172 (n=1) | 1.6 (n=1) | 20-31d at 18°C | | | | | | | | Huff&McCafferty (1974) |
| Heptageniidae | *Rhithrogena hybrida* | *Rhrithrogena cf. hybrida* | West PAL | Austria | | Seebach | **Sexual** | tycho. | both sexes | up to 2300 | 0.1 (0-1.6, n=22) | 78-294d at 5-8°C; 37-72d at 10-15°C | | artificial fertilisatic | 70-142d at 4.5-7°C; 14-54d at 10-20°C | | | | | Humpesch (1980)b; Humpesch&Elliott (1980); Humpesch (1982) |
| Heptageniidae | *Rhithrogena loyolaea* | | West PAL | Austria | | Herrnalmbach | **Sexual** | tycho. | both sexes | | 1.8 (0-9.2, n=7) | 175-340d at 4-6.5°C | | artificial fertilisatic | 170-340d at 5°C | | | | | Humpesch (1980)b; Humpesch&Elliott (1980) |
| Heptageniidae | *Rhithrogena picteti* | *Rhithrogena sp.* | West PAL | France | Isère | La Drevenne, à Saint | **Sexual** | tycho. | both sexes | 1276-1806 (n=2) | 3.1 (0.2-5.1, n=2) | | | | | | | | | Degrange (1960) |
| Heptageniidae | *Rhithrogena picteti* | *Rhithrogena sp.* | West PAL | France | Isère | Le Doménon, à Reve | **Sexual** | tycho. | both sexes | | 0.2 | | | | | | | | | Degrange (1960) |
| Heptageniidae | *Rhithrogena picteti* | *Rhithrogena sp.* | West PAL | France | Haute-Savoie | Le Fier, au pont de N | **Sexual** | tycho. | both sexes | | 0.8 | | | | | | | | | Degrange (1960) |
| Heptageniidae | *Rhithrogena picteti* | | West PAL | Switzerland | Vaud | 46.82108, 6.62107 | **Sexual** | tycho. | female-biased: 0.6 (n=400) | | 0.3 (0-0.5, n=3) | | | | | | | | | Maud Liégeois (personal observations) |
| Heptageniidae | *Rhithrogena semicolorata* | *Rhithrogena semitincta* | West PAL | France | Isère | Le Lanfrey, au Fonta | **Mixed (sympatry)** | fac. (mixed in symp | both sexes | 2576.5 (2465-27 | 14.9 (10-20, n=2) | 23-63d | | | | | | | | Degrange (1960) |
| Heptageniidae | *Rhithrogena semicolorata* | *Rhithrogena semitincta* | West PAL | France | Isère | Ruisseau de St-Etien | **Mixed (sympatry)** | tycho. | both sexes | 3313-3581 (n=2) | 3.4 (n=1) | 23-63d | | | | | | | | Degrange (1960) |
| Heptageniidae | *Rhithrogena semicolorata* | *Rhithrogena semitincta* | West PAL | France | Isère | Ruisseau de Lalley | **Mixed (sympatry)** | sex.? | both sexes | 1000-2000 | | | | | 17d at 18°C | 48 hours | | | | Degrange&Perrier (1957); Degrange (1960) |
| Heptageniidae | *Rhithrogena semicolorata* | | West PAL | England | | Wilfin Beck | **Mixed (sympatry)** | sex.? | both sexes | | | | | up to 46 | 51-123d at 6-8°C; 7-69d at 12-20°C | | | | "mated females colle | Humpesch&Elliott (1980) |
| Heptageniidae | *Rhithrogena semicolorata* | | West PAL | England | | River Lune | **Mixed (sympatry)** | sex.? | both sexes | | | | | artificial fertilisatic | 10-57d at 8-21°C | | | | | Humpesch&Elliott (1980) |
| Heptageniidae | *Rhithrogena semicolorata* | | West PAL | Czech Republic | Čertova Stěna | Kleštínský brook | **Mixed (sympatry)** | | both sexes | | | | | | | | | 18+XX:XY | | Soldan&Putz (2000) |
| Heptageniidae | *Stenacron interpunctatum* | *Stenonema interpunctatum* | NEA | Indiana | | | **Sexual** | tycho. | both sexes | | <0.1 | 32d at 18°C | | | | | | | | Huff&McCafferty (1974) |
| Heptageniidae | *Stenacron interpunctatum* | *Stenacron interpunctatum fr* | NEA | Maine | Aroostook Co. | Logan Brook | **Sexual** | tycho. | balanced | 974 | 9.1 (n=1); dvpmt: 15 | 21.6d (17-47 at 22±2°C) | | | | | | | | Mingo (1978) |
| Heptageniidae | *Stenonema femoratum* | | NEA | Indiana | | | **Mixed (sympatry)** | mixed (sympatry) | balanced: ~1:1 | 3810.7 (n=20) | 8.3 (0.5-36.5, n=18) | 18-29d at 18°C | | forced copulation: | 12-15d at 18°C | | | | | Huff&McCafferty (1974); McCafferty&Huff (1974) |
| Heptageniidae | *Stenonema femoratum* | | NEA | Ontario | Haliburton Co. | Thompsons Creek | **Mixed (sympatry)** | sex. | both sexes | | | | | forced copulation: | 7-12d | | | | | Rowe et al. (1988) |
| Heptageniidae | *Stenonema femoratum* | | NEA | Missouri | Phelps Co. | Beaver Creek (BC): 3 | **Mixed (sympatry)** | mixed (sympatry) | female-biased: 0.81 (n=21, 2years) | | 7.3 (0-53.2, n=15) | | | | | | | | | Ball (2002) |
| Heptageniidae | *Stenonema femoratum* | | NEA | Missouri | Cole Co. | Honey Creek (HNC): | **Mixed (sympatry)** | mixed (sympatry) | female-biased: 0.76 (n=74, 2years) | | 4.3 (0-25.5, n=11) | | | | | | | | | Ball (2002) |
| Heptageniidae | *Stenonema femoratum* | | NEA | Missouri | Boone Co. | Rock Bridge Gans Cr | **Mixed (sympatry)** | mixed (sympatry) | female-biased: 0.70 (n=47, 3 years) | | 5.7 (0-77.9, n=38) | ~12d at 23-25°C | | | ~7d at 23-25°C | | automixis (both homozygous and heterozygous offsprin | | | Ball (2001, 2002) |
| Heptageniidae | *Stenonema femoratum* | | NEA | Missouri | Callaway Co. | Cedar Creek (CC): 38 | **Mixed (sympatry)** | mixed (sympatry) | female-biased: 0.66 (n=158, 2years) | | 6.6 (0-46.5, n=64) | | | | | | | | | Ball (2002) |
| Heptageniidae | *Stenonema femoratum* | | NEA | Missouri | Boone Co. | Grindstone Creek (G | **Mixed (sympatry)** | mixed (sympatry) | female-biased: 0.69 (n=140, 2years) | | 8.6 (0-57.3, n=19) | | | | | | | | | Ball (2002) |
| Heptageniidae | *Stenonema femoratum* | | NEA | Missouri | Shannon Co. | Jacks Fork River (JF): | **Mixed (sympatry)** | mixed (sympatry) | female-biased: 0.78 (n=36, 2years) | | 5.3 (0-31, n=28) | | | | | | | | | Ball (2002) |
| Heptageniidae | *Stenonema femoratum* | | NEA | Missouri | Crawford Co. | Meramec River (MR) | **Mixed (sympatry)** | mixed (sympatry) | female-biased: 0.71 (n=48, 2years) | | 5.5 (0-45.8, n=45) | | | | | | | | | Ball (2002) |

| | | | | | | | | | | | | | | | | | | | | |
|---|---|---|---|---|---|---|---|---|---|---|---|---|---|---|---|---|---|---|---|---|
| Heptageniidae | *Stenonema femoratum* | | NEA | Missouri | Callaway Co. | Little Dixie Lake (LD) | **Mixed (sympatry)** | fac. (mixed in symp | female-biased: 0.71 (n=439, 3years) | | 14.5 (0-67.7, n=71) | ~12d at 23-25°C | | | ~7d at 23-25°C | | automixis (both homozygous and heterozygous offsprin | | | Ball (2001, 2002) |
| Heptageniidae | *Stenonema femoratum* | | NEA | Missouri | Phelps Co. | Little Prairie Lake (LP | **Mixed (sympatry)** | mixed (sympatry) | female-biased: 0.68 (n=114, 2years) | | 6.9 (0-32.4, n=36) | | | | | | | | | Ball (2002) |
| Heptageniidae | *Stenonema femoratum* | | NEA | Missouri | Boone Co. | Rocky Forks Lake (RF | **Mixed (sympatry)** | mixed (sympatry) | female-biased: 0.73 (n=256, 2years) | | 3.3 (0-16.1, n=43) | | | | | | | | | Ball (2002) |
| Heptageniidae | *Stenonema femoratum* | | NEA | Missouri | Cole Co. | Winegar Lake (WL): | **Mixed (sympatry)** | fac. (mixed in symp | female-biased: 0.74 (n=205, 2years) | | 10.7 (0-41.6, n=5) | | | | | | | | | Ball (2002) |
| Heptageniidae | *Stenonema femoratum* | | NEA | Missouri | Callaway Co. | Whetstone Lake (WS | **Mixed (sympatry)** | mixed (sympatry) | female-biased: 0.70 (n=324, 2years) | | 7.4 (0-63.4, n=57) | | | | | | | | | Ball (2002) |
| Isonychiidae | *Isonychia bicolor* | | NEA | Pennsylvania | Chester Co. | WCC: 39°53'N, 75°47 | **Sexual** | tycho. | both sexes | 732-3302 | <0.1 (0-0.1) | | | artificial fertilisatic | 25-54d at 17°C | 80% within a week, 20% after 4-5 weeks | | | | Sweeney (1978); Sweeney&Vannote (1987) |
| Leptophlebiidae | *Choroterpes picteti* | | West PAL | France | Ardèche | L'Eyrieux, en amont | **Sexual** | sex. | both sexes | 724 | 0 | | | | | | | | females mate, lay and | Degrange (1960) |
| Leptophlebiidae | *Choroterpes sp.* | | ORI | China | Hong Kong | | **Mixed (sympatry)** | fac. (mixed in symp | both sexes | 663 (637-703, n= | 13 ±5.3 (n=3) | 14.6 ±0.8d (11-25, at 22±2°C) | | | | | | | | Salas&Dudgeon (1999) |
| Leptophlebiidae | *Fittkaulus cuiabae* | | NEO | Brazil | Mato Grosso | | **"Obligate"** | obl. | females only | | | | | | | | | | males are rares | Savage (1986); Frederico F. Salles (personal communication) |
| Leptophlebiidae | *Fittkaulus cururuensis* | | NEO | Brazil | Pará, Espírito Santo | | **"Obligate"** | obl. | females only | | | | | | | | | | males are rares | Savage (1986); Frederico F. Salles (personal communication) |
| Leptophlebiidae | *Habroleptoides auberti* | *Habrophlebia auberti* | West PAL | France | Haute-Savoie | Le Fier, au pont de N | **Sexual** | sex. | both sexes | 1412-1685 | 0 | | | | | | | | | Degrange (1960) |
| Leptophlebiidae | *Habroleptoides modesta* | *Habrophlebia modesta* | West PAL | France | Isère | Le Charmeyran, à La | **Sexual** | tycho. | both sexes | 932-2324 | 0.01 (0-0.04, n=7) | 49d at 11-15°C | | natural fertilisatior | 16-27d at 11-14°C | | | | mated females catch | Degrange (1960) |
| Leptophlebiidae | *Habrophlebia lauta* | | West PAL | France | Isère | Le Méaudret, à Méa | **Sexual** | sex. | both sexes | 450-790 | 0 | | | | | | | | | Degrange (1960) |
| Leptophlebiidae | *Leptophlebia cupida* | | NEA | Alberta | | Bigoray River: 53°31 | **Sexual** | sex. | both sexes | 2959 | 0 | | | artificial fertilisatic | 10-43d at 20°C | mostly within 10-14 days | | | | Clifford et al. (1979) |
| Leptophlebiidae | *Leptophlebia cupida* | | NEA | Ontario | Haliburton Co. | Plastic Lake | **Sexual** | sex.? | both sexes | | | | | natural fertilisatior | 8-17d | | | | mated females catch | Rowe et al. (1988) |
| Leptophlebiidae | *Leptophlebia cupida* | | NEA | Eastern North America | | | **Sexual** | tycho. (mixed) | both sexes | | 13 | | | | | | | | | Sweeney&Vannote (1987) |
| Leptophlebiidae | *Leptophlebia marginata* | | West PAL | France | Drôme | Le Rhône, à Valence | **Sexual** | sex. | both sexes | | 0 | | | | | | | | | Degrange (1960) |
| Leptophlebiidae | *Leptophlebia vespertina* | | West PAL | France | Savoie | Lac de Saint-André | **Sexual** | tycho. | both sexes | 1016-1612 | 2.2 (n=1) | 42-51d at 13-17°C | | | | | | | | Degrange (1960) |
| Leptophlebiidae | *Leptophlebia vespertina* | | West PAL | Wales | | Llyn Dinas Lake | **Sexual** | | both sexes | | | | | | 21-26d at 14.5-20.5°C | 90% within 48h | | | | Brittain (1976) |
| Leptophlebiidae | *Paraleptophlebia debilis* | | NEA | Eastern US | | | **Sexual** | tycho.? | both sexes | | hatching | | | | | | | | "species that we had | Funk et al. (2010) |
| Leptophlebiidae | *Paraleptophlebia submarginata* | | West PAL | France | Savoie, Isère | | **Sexual** | sex. | both sexes | | 0 | | | | | | | | | Degrange (1960) |
| Leptophlebiidae | *Paraleptophlebia submarginata* | | West PAL | Czech Republic | Holubov | Křemžský brook | **Sexual** | | both sexes | | | | | | | | | (12-14)+XX:XY | | Soldan&Putz (2000) |
| Leptophlebiidae | *Thraulodes sp.* | | NEO | Costa Rica | Rìo Tempisquito | 10°57'25''N, 85°29'4 | **"Obligate"** | obl. | females only: 1.00 | | (74.2-95.2, n=3) | median: 19 ±1d at 20±1°C | | | | | | | thelytokous parthenc | Jackson&Sweeney (1995) |
| Leptophlebiidae | *Thraulus bellus* | | West PAL | France | Isère | Lac de Laffrey | **Sexual** | sex. | both sexes | 1371 | 0 | | | | | | | | females mate and lay | Degrange (1960) |
| Oligoneuriidae | *Oligoneuriella rhenana* | | West PAL | France | Ardèche | L'Eyrieux, en amont | **Sexual** | sex. | both sexes | | 0 | | | | | | | | females mate at subi | Degrange (1960) |
| Oligoneuriidae | *Oligoneuriella rhenana* | | West PAL | Czech Republic | Kaplice | Malše River | **Sexual** | | both sexes | | | | | | | | | 14+XX:XY | | Soldan&Putz (2000) |
| Palingeniidae | *Palingenia longicauda* | | West PAL | Hungary | | Tisza River | **Facultative** | fac. | balanced: ~1:1 (n=300) | | ~52 | 11-46d at 20°C | | artificial fertilisatic | 4-39d at 20°C | | | | /!\% of developemer | Landolt et al. (1997) |
| Polymitarcyidae | *Ephoron album* | | NEA | North America | | Lake Erie | **Sexual** | tycho. | balanced: 0.47 (n=559) | 908 (593-1672, n | (8-10) | diapause (min. 200d) | | much higher than unfertilised | | | | | females mate, lay and | Britt (1953, 1962) |
| Polymitarcyidae | *Ephoron eophilum* | | East PAL | Japan | Ibaraki Prefecture: Tsu | Sakura-gawa River: 3 | **Mixed (sympatry)** | sex. | both sexes | | | | | 97.4 (78.1-99.7, n=16) | | | automixis: eit | 14+XX:XO | /!\% of developemer | Tojo et al. (2006); Sekine et al. (2015)b |
| Polymitarcyidae | *Ephoron eophilum* | | East PAL | Japan | Ibaraki Prefecture: Jos | Kinu-gawa River: 36° | **Mixed (sympatry)** | fac. (mixed in symp | both sexes | 352-1863 | 15.5 (0-70.2, n=7) | | | | | | automixis: eit | 14+XX:XO | /!\% of developemer | Sekine et al. (2013); Sekine et al. (2015)b |
| Polymitarcyidae | *Ephoron shigae* | | East PAL | Japan | Okayama Prefecture: C | Asahi-gawa River: 34 | **Mixed** | tycho. | balanced: 0.5 | | 0.3 (0-8, n=25) | | | 98.4 ±0.7 (97.1-99. | 134-147d (diapause) at 2( | 10-12 days | | | females mate, lay and | Watanabe&Takao (1991); Tojo et al. (2006); Sekine&Tojo (2010)b |
| Polymitarcyidae | *Ephoron shigae* | | East PAL | Japan | Tochigi Prefecture: Uts | Kinu-gawa River: 36° | **Mixed** | obl. | females only: 1.00 | | > 80 | | | | | | | | | Nakamura (1985, in japanese) |
| Polymitarcyidae | *Ephoron shigae* | | East PAL | Japan | Nagaro Prefecture: Chi | Chikuma-gawa River | **Mixed** | obl. | females only: 1.00 | | 89.2 ±4.6 (76.2-96.9, n=23) | | | | | | | females: 10+XX | | Tojo et al. (2006); Sekine&Tojo (2010)b |
| Polymitarcyidae | *Ephoron shigae* | | East PAL | Japan | Ibaraki Prefecture: Tsu | Sakura-gawa River: 3 | **Mixed** | obl. | females only: 1.00 | | 84.2 ±2.0 (81.1-87.4, n=20) | | | | | | automixis with terminal fusion | | diploid telytokous; pr | Tojo et al. (2006); Sekine&Tojo (2010)a |
| Polymitarcyidae | *Ephoron shigae* | | East PAL | Japan | Aichi Prefecture: Nago | Shonai-gawa River: 3 | **Mixed** | mostly obl. | almost females only: 0.99 | | 78.5 ±6.0 (69.3-86.8, n=13) | | | | | | | | | Tojo et al. (2006) |
| Polymitarcyidae | *Ephoron shigae* | | East PAL | Japan | Tokyo Prefecture: Hino | Hino-yosui Canal: 36 | **Mixed** | fac. (mixed in symp | balanced | | 14.1 ±13.5 (0.3-34.7, n=10) | | | 98.1 ±1.9 (94.8-99.9, n=6) | | | | 10+XX:XO | | Sekine&Tojo (2010)b |
| Polymitarcyidae | *Ephoron shigae* | | East PAL | Japan | Saitama Prefecture: Ko | Ara-kawa River: 36°5 | **Mixed** | fac. (mixed in symp | balanced | | 21.5 ±27.5 (0-53.3, n=5) | | | | | | | | | Sekine&Tojo (2010)b |
| Polymitarcyidae | *Ephoron virgo* | | West PAL | France | Indre-et-Loire | La Vienne, à Chinon | **Sexual** | sex. | both sexes | | 0 | | | | diapause | | | | | Degrange (1960) |
| Polymitarcyidae | *Ephoron virgo* | | West PAL | Germany | | several populations | **Sexual** | sex. | balanced: 0.49 (n=2895, 2 | 3194 (2333-4058, n=172) | | | | > 95 | diapause | within a month | | | | Kureck et al. (2014) |
| Potamanthidae | *Potamanthus luteus* | | PAL (west + east) | France | Rhône | La Saône, à Belleville | **Sexual** | sex. | both sexes | | 0 | | | | | | | | | Degrange (1960) |
| Potamanthidae | *Potamanthus luteus* | | PAL (west + east) | Czech Republic | Skryje | Berounka River | **Sexual** | | both sexes | | | | | | | | | 6+XX:XO | | Soldan&Putz (2000) |
| Prosopistomatidae | *Prosopistoma foliaceum* | *Binoculus pennigerus* | West PAL | France | Drôme | Le Rhône, à Valence | **Sexual** | sex. | both sexes | 174 | 0 | | | | | | | | | Degrange (1960) |
| Prosopistomatidae | *Prosopistoma pearsonorum* | | AUS | Australia | north Queensland | | **"Obligate"** | obl. | females only | | | | | | | | | | | Campbell&Hubbard (1998); Ian Campbell (personal communication) |
| Siphlonuridae | *Siphlonisca aerodromia* | | NEA | Maine | Washington Co. | Tomah Stream: 45°2 | **Facultative** | fac. | female-biased: 0.63 (4 ye | (313-559, n=16) | 54.8 ±8.1 (34.9-85.6, n=6); dvpmt: 62.9 ±9.2 (46.9-92.1) | | | | diapause | | | | | Gibbs&Mingo (1986); Gibbs&Siebenmann (1996) |
| Siphlonuridae | *Siphlonurus aestivalis* | | West PAL | France | Isère | Ruisseau affluent du | **Sexual** | tycho. | both sexes | 933-2678 | 5.4 | 159-175d at 15°C (diapause) | | | | | | | | Degrange (1954, 1960) |
| Siphlonuridae | *Siphlonurus lacustris* | | West PAL | France | Haute-Savoie | Le Fier, au Pont de N | **Mixed (sympatry)** | fac. (mixed in symp | both sexes | 1712-2499 | 11.9 (0-33, n=9) | 40-163d at 18°C | | forced copulation (failed) | | | | | | Degrange (1955, 1960) |
| Siphlonuridae | *Siphlonurus lacustris* | | West PAL | Czech Republic | Modrava | Modravský brook | **Mixed (sympatry)** | | both sexes | | | | | | | | | 16+XX:XY | | Soldan&Putz (2000) |
